# Structural insights into the complex of oncogenic KRas4B^G12V^ and Rgl2, a RalA/B activator

**DOI:** 10.1101/2022.10.10.511529

**Authors:** Mishal Tariq, Teppei Ikeya, Naoyuki Togashi, Louise Fairall, Syun Kamei, Sannojah Mayooramurugan, Lauren R. Abbott, Anab Hasan, Carlos Bueno-Alejo, Sakura Sukegawa, Beatriz Romartinez-Alonso, Miguel Angel Muro Campillo, Andrew J. Hudson, Yutaka Ito, John W.R. Schwabe, Cyril Dominguez, Kayoko Tanaka

**Author notes:** Corresponding author: *Kayoko Tanaka **Email:**.

## Abstract

About a quarter of total human cancers carry mutations in Ras isoforms. Accumulating evidence suggests that small GTPases, RalA and RalB, and their activators, Ral guanine nucleotide exchange factors (RalGEFs), play an essential role in oncogenic Ras-induced signalling. We studied the interaction between human KRas4B and the Ras association (RA) domain of Rgl2 (Rgl2^RA^), one of the RA-containing RalGEFs. We show that the G12V oncogenic KRas4B mutation changes the interaction kinetics with Rgl2^RA^. The crystal structure of the KRas4B^G12V^: Rgl2^RA^ complex shows a 2:2 heterotetramer where the Switch I and Switch II regions of each KRas^G12V^ interact with both Rgl2^RA^ molecules. This structural arrangement is highly similar to the HRas^E31K^:RALGDS^RA^ crystal structure and is distinct from the well-characterised Ras:Raf complex. Interestingly, the G12V mutation was found at the dimer interface of KRas4B^G12V^ with its partner. Our study reveals a potentially distinct mode of Ras:effector complex formation by RalGEFs, and offers a possible mechanistic explanation for how the oncogenic KRas4B^G12V^ hyperactivates the RalA/B pathway.

## Summary Blurb

The work reports the KRas4B.G12V oncogenic mutation alters the binding kinetics with Rgl2, a RalA/B activator, and the V12 resides at the KRas4B(G12V)•Rgl2 complex interface in the crystal structure.

## Introduction

Ras belongs to a family of small G protein that switches between two states, GDP-bound form (Ras^GDP^) and GTP-bound form (Ras^GTP^), and regulates a wide range of cellular activities (Cox and Der, 2010). Guanine nucleotide exchange factors (GEFs), typically activated by growth factor signalling, mediate the conversion of Ras^GDP^ to Ras^GTP^, whilst the GTP-bound status lasts only transiently as the intrinsic GTPase activity, aided by the GTPase activating Proteins (GAPs), hydrolyses the bound GTP into GDP (Vetter and Wittinghofer, 2001). Ras acts as a signalling hub where the Ras^GTP^, but not Ras^GDP^, physically interacts with multiple Ras effectors, which then transmit the signal to downstream molecules, including ERK, Akt and small G proteins RalA and RalB (Simanshu et al., 2017).

There are three human *RAS* genes, *KRAS*, *NRAS* and *HRAS*, and as *KRAS* is alternatively spliced at exon 4, the three *RAS* genes produce four Ras isoforms KRas4A, KRas4B, NRas and HRas (Castellano and Santos, 2011). Among them, *KRAS4B* typically represents more than half of all *RAS* transcripts (Newlaczyl et al., 2017). The Catalogue Of Somatic Mutations In Cancer (COSMIC) shows that about 25% of all cancers carry mutations in *RAS* genes, and *KRAS* is responsible for about 70% of these mutations (COSMIC, v.95). Hence, it is vital to obtain more insights into how the oncogenic KRas signalling leads to cancers.

Extensive earlier biochemical studies revealed that oncogenic *RAS* mutations cause a reduction of the GTP hydrolysis rate, generating an increased population of Ras^GTP^ (Bollag and McCormick, 1991; Der et al., 1986; Gibbs et al., 1984; Manne et al., 1985; McGrath et al., 1984; Scheffzek et al., 1997). This likely leads to an overactivation of the downstream signalling pathways and therefore is considered the major cause of the KRas oncogenicity. In addition, recent biochemical, structural and molecular modelling studies indicate that oncogenic *RAS* mutations affect the interaction kinetics with the effector molecules and may trigger a biased overactivation of a set of effectors (Hunter et al., 2015; Mazhab-Jafari et al., 2015; Pantsar et al., 2018; Smith and Ikura, 2014). Therefore, obtaining more insights into the Ras-effector interaction mechanisms is essential to understanding the oncogenic-Ras mediated tumorigenesis process.

Effector molecules that have been shown to interact with Ras all carry a domain consisting of a ubiquitin-fold structure, although the primary sequences of these domains show a limited homology (Kiel and Serrano, 2006). They are classified into three categories according to Uniprot PROSITE-ProRule Annotation based on their primary sequences; Ras-associating (RA) (PRU00166), Ras-binding domain (RBD) (PRU002262), and phosphatidylinositol 3-kinase Ras-binding domain (PI3K-RBD) (PRU00879). In this study, we follow these annotations.

Among Ras effectors, Raf kinase and PI3K, which triggers ERK and Akt signalling, respectively, have attracted much attention, especially because they were found not only to be activated by oncogenic Ras but also to be able to carry oncogenic mutations themselves (Chalhoub and Baker, 2009; Maurer et al., 2011). However, accumulating evidence suggests that the misregulation of Ral GTPases, RalA and RalB, rather than ERK and Akt signalling pathways, may act as the initial trigger of oncogenic-Ras induced tumorigenesis. For example, in the oncogenic K-ras(G12D) knock-in mouse model, although neoplastic and developmental defects were observed, hyper-activation of ERK or Akt was undetected (Tuveson et al., 2004). In humans, ERK hyper-activation is often missing in cancer cell lines and tissues with oncogenic Ras mutations, whereas RalA and RalB are essential in oncogenic-Ras induced cell proliferation, motility and tumorigenesis (Campbell et al., 2007; Lim et al., 2005; Lim et al., 2006; Luo and Sharif, 1999; Miller et al., 1997; Yip-Schneider et al., 1999; Yip-Schneider et al., 2001; Zago et al., 2018). Therefore, a biased misregulation of the Ral GTPases may be the critical feature behind Ras oncogenicity. Indeed, NMR-based effector competition assays suggested the intriguing possibility that the oncogenic Ras molecules may develop an altered effector preference leading to a biased hyperactivation of RalA (Smith and Ikura, 2014).

Activation of RalA and RalB is mediated by guanine nucleotide exchange factors (RalGEFs). There are eight RalGEFs reported for humans, and four of them, RALGDS, Rgl1, Rgl2 and Rgl3, have a Ras-association (RA) domain responsible for Ras-binding (Apken and Oeckinghaus, 2021) RALGDS was the first to be identified as a Ras effector among these RalGEFs and is by far the most studied (Neel et al., 2011). RALGDS is one of the first effector molecules that was crystallised together with an active Ras; rat RALGDS was co-crystalised with human HRas harbouring an E31K mutation, which helped complex formation (Huang et al., 1998). Meanwhile, a recent modelling approach, integrating proteomic data of Ras, its 56 effectors and their interaction affinities, predicts that Rgl2 would generate the highest concentration of Ras:effector complex among RalGEFs in 28 out of 29 healthy human tissues (Catozzi et al., 2021). Furthermore, Rgl2 plays a critical role in oncogenic-Ras induced tumour phenotypes (Vigil et al., 2010), and overexpression of the Rgl2^RA^ interferes with Ras-mediated signaling, likely by competitively titrating active Ras molecules (Fischer et al., 2003; Peterson et al., 1996).

Human Rgl2 and its mouse homologue, Rlf, were initially identified through yeast two-hybrid screenings and were shown to interact with Ras through the RA domain *in vitro* (Esser et al., 1998; Ferro et al., 2008; O’Gara M et al., 1997; Peterson *et al*., 1996; Wolthuis et al., 1996). Furthermore, *in vivo* interaction of Rgl2 and Ras was reported in a BioID-based proteomics study where Rgl2 was among other proximal interactors identified using HRas.G12V expressed in bladder cancer cells (Kovalski et al., 2019). In addition, the full-length Rlf was shown to interact with HRas.G12V and acted as a RalGEF when expressed in COS-7 cells (Wolthuis et al., 1997). However, the Ras:Rgl2 complex interface at an atomic level has not been explored. Furthermore, whether oncogenic mutations influence the interaction kinetics or not is unclear, making it difficult to appreciate the impact of an oncogenic mutation in activating the RalGEF signalling branch that plays an essential role in the oncogenic Ras-mediated tumorigenesis.

To address the question, we examined the mode of interaction between human KRas4B and the RA of Rgl2 (Rgl2^RA^) in this report. We observed a change in the interaction kinetics between KRas4B and the Rgl2^RA^ upon the introduction of the G12V oncogenic mutation. Our crystal structure of KRas4B^G12V^:Rgl2^RA^ complex shows a heterotetramer formation, highly similar to the reported HRas^E31K^:RALGDS^RA^ complex, but distinct from other Ras:effector complexes, including Ras:Raf1. Interestingly, in our structure, the G12V oncogenic mutation is located at the dimer interface of KRas4B^G12V^ with its homodimeric partner. Our findings provide an interesting possibility that KRas4B^G12V^ oncogenicity might be contributed by the altered interaction kinetics with Rgl2.

## Results

### The G12V oncogenic mutation causes a change in the interaction kinetics between the KRas4B and the Rgl2^RA^

Throughout this study, we used recombinant bacteria constructs of human KRas4B lacking the C-terminal hyper-variable region and the Rgl2^RA^ consisting of amino acids 643 – 740 of human Rgl2 (Supplementary Fig. S1A). The binding of Rgl2^RA^ and KRas4B^G12V^ was confirmed for the active KRas4B^G12V^ loaded with a non-hydrolysable GTP analogue (Guanosine-5’-[(β,γ)- imido]triphosphate (GMPPNP)), but not with the GDP-loaded KRas4B^G12V^, by GST-pulldown assays (Supplementary Fig. S1B). To examine whether the binding mode of Rgl2^RA^ differs between KRas4B^WT^ and KRas4B^G12V^, the binding kinetics were measured by biolayer interferometry (BLI). We noticed that both KRas4B^WT^ and KRas4B^G12V^, purified at 4°C at all times, retained GTP as the major bound guanine nucleotide. In addition, in our hands, the efficacy of GMPPNP loading could vary between samples, whilst the efficacy of GTP loading was highly reproducible. We also confirmed that the KRas4B^WT^ and KRas4B^G12V^ samples that were subjected to the incubation condition of the BLI assay (20°C for 30 min) were still associated with GTP (Fig. 1A). Therefore, we loaded the samples with GTP and used these GTP-bound samples for the BLI experiment so that we could minimize possible artefacts caused by GMPPNP.

**Figure 1.**
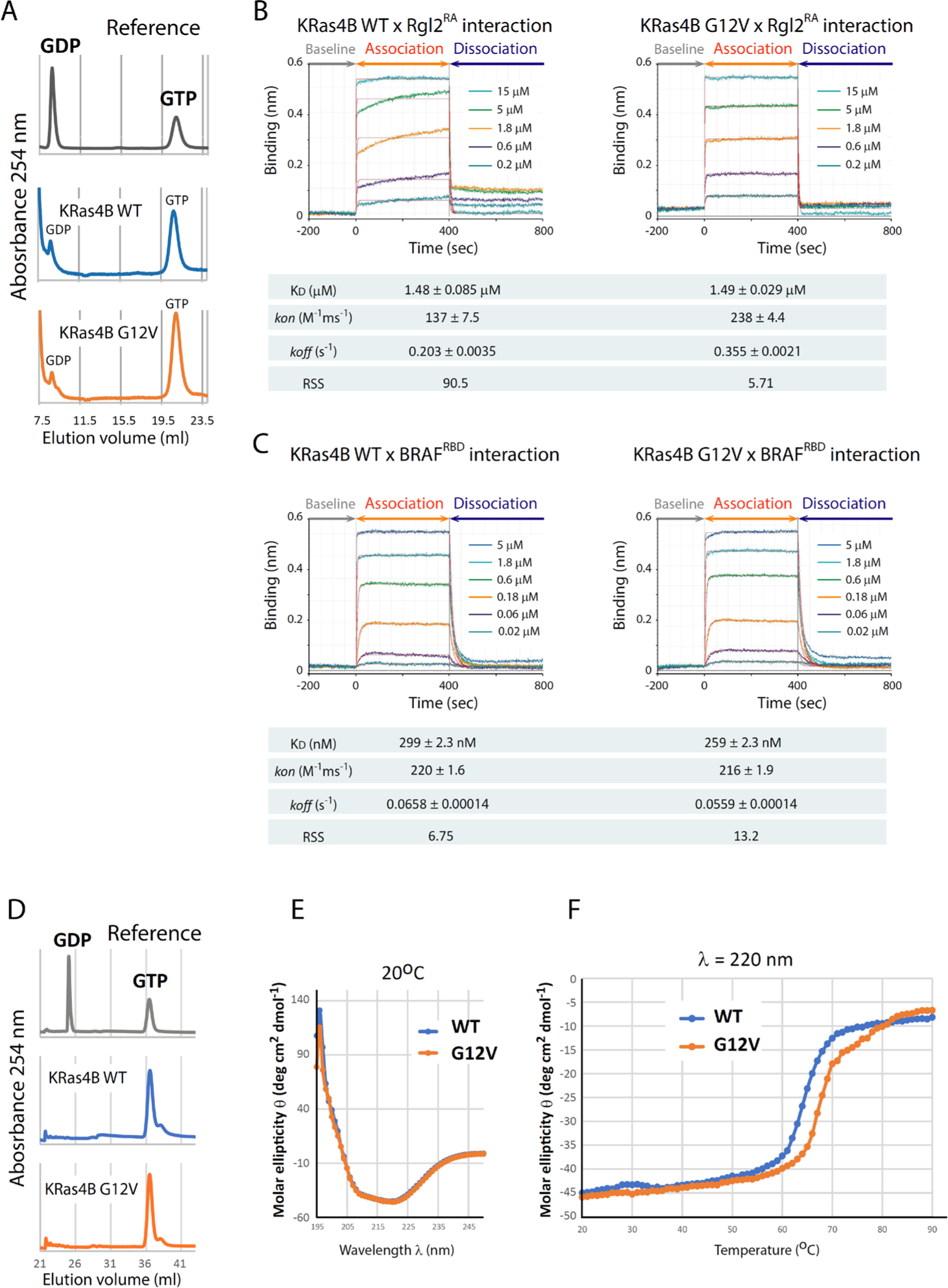
KRas4B^G12V^ exhibits a different binding kinetics towards Rgl2^RA^ than KRas4B^WT^ and is structurally more stable. (A and D) KRas4B^WT^ and KRas4B^G12V^ samples used for biolayer Interferometry (BLI) and Circular Dichroism (CD) were confirmed to be loaded with GTP. The nucleotide-binding status of KRas4B^WT^ and KRas4B^G12V^ were examined by denaturating the proteins and detecting the released nucleotides by anion exchange chromatography. Samples of pure GDP or GTP were used as references. (B) KRas4B^WT^ and KRas4B^G12V^ show different binding kinetics towards Rgl2^RA^. BLI was used to measure the binding kinetics of KRas4B (analyte) to immobilized GST-Rgl2^RA^ (ligand). GST-Rgl2^RA^ was immobilized on the bio-sensors, and varying concentrations of free KRas4B^WT^ (left panel) and KRas4B^G12V^ (right panel) were provided, and the interactions were measured at 20°C. The 1:1 binding model was fitted to the data using Octet Analysis Studio 13.0 (Sartorius). The fitted curves are shown in red. The deduced KD values of these two cases are similar: approximately 1.48 µM (WT) and 1.49 µM (G12V). Meanwhile, the *kon* values are approximately 137 (M^-1^ms^-1^) (WT) and 238 (M^-1^ms^-1^) (G12V), and *koff* values are approximately 0.20 (s^-1^) (WT) and 0.36 (s^-1^) (G12V), indicating that the G12V mutation may cause the interaction more dynamic. The high residual sum of squares (RSS) value (90.5) for the wildtype case indicates that the 1:1 binding model fitting for the wildtype case is not as good as for the G12V case. (C) KRas4B^WT^ and KRas4B^G12V^ show highly similar binding kinetics towards BRAF^RBD^. BLI was conducted in the same manner as (B), except using GST-BRAF^RBD^ as the ligand. The 1:1 binding model fitting (shown in red) predicts the KD values of these two cases to be about 299 nM (WT) and 259 nM (G12V), respectively. The *kon* and *koff* values are comparable between the two cases, and the RSS values ensure the 1:1 binding model fitting is suitable for both the wildtype and the G12V cases. (D) CD spectra of KRas4B^WT^ and KRas4B^G12V^ (20 μM) at 20°C. (E) CD signal intensity at 220 nm as a function of temperature from 20°C to 90°C.

The sensorgram curves representing the on/off kinetics between KRas4B and Rgl2^RA^ were distinct when using the KRas4B^WT^ or KRas4B^G12V^ proteins (Fig. 1B). When the KRas4B-Rgl2^RA^ binding kinetics results were model-fitted using a 1:1 binding model, the *kon* and *koff* values were larger in the case of the G12V mutant, indicating more dynamic interaction between the G12V mutant and Rgl2^RA^ (Fig. 1B). Meanwhile, the KD values were comparable between the KRas4B^WT^ (about 1.48 µM) and KRas4B^G12V^ (about 1.49 µM)(Fig. 1B), and the elution profiles of the size exclusion chromatography showed little difference between the wildtype and the G12V mutant (Supplementary Fig. S1, D and E).

Interestingly, the KRas4B^WT^-Rgl2^RA^ sensorgram curve was more consistent with a 2:1 heterogeneous binding model fitting or a 1:2 bivalent binding model fitting (Data Analysis HT Software, Sartorius), as indicated by the improvement of the residual sum of squares (RSS) value; from about 90.5 (the 1:1 model, Fig. 2B) to about 1.05 (the 2:1 model, Supplementary Fig. S2A) or 1.85 (the 1:2 model, Supplementary Fig. S2A). Therefore, the KRas4B^WT^-Rgl2^RA^ interaction does not correspond to a simple 1:1 binding mode. In contrast, for the KRas4B^G12V^- Rgl2^RA^ sensorgram curve, the RSS values for the 1:1 and the 2:1 model fittings were comparable; about 5.71 for the 1:1 model (Fig. 1B) and about 4.40 for the 2:1 model (supplementary Fig. S2B). Furthermore, in the 2:1 heterogeneous model, “Binding type 2” represents 95-100% of the whole population, where the values for the KD, *kon* and *koff* are all similar to the ones deduced from the 1:1 binding model (Supplementary Fig. S2B). These results strongly indicate that the binding of the KRas4B^G12V^-Rgl2^RA^ complex is compatible with a 1:1 binding.

**Figure 2.**
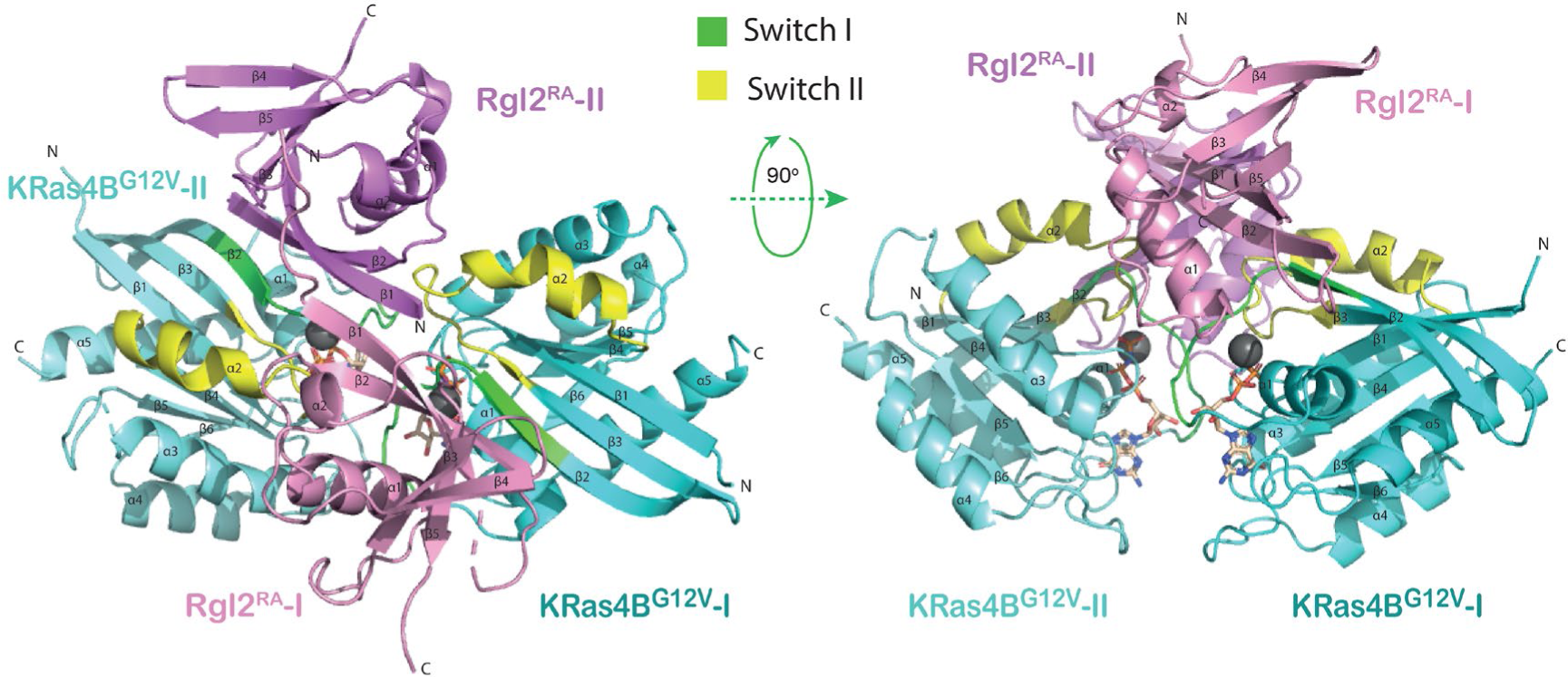
Crystal structure of the KRas4B^G12V^:Rgl2^RA^ 2:2 heterotetramer. Cartoon representation of the structure of the heterotetramer complex of KRas4B^G12V^ and Rgl2^RA^ with top view and side view; the two KRas4B^G12V^ molecules are shown in dark and pale cyan and the two molecules of Rgl2^RA^ in pink and violet. Switch-I and Switch-II regions of KRas4B^G12V^ are shown in green and yellow, respectively, and the α-helix and β-sheets are numbered for each chain. The Mg^2+^ is shown as a grey sphere. The structure shows that each KRas4B^G12V^ molecule interacts with two Rgl2^RA^ molecules (referred to as I and II) at Switch-I and Switch-II individually.

The difference in kinetics of binding observed for KRas4B^WT^ and KRas4B^G12V^ mutant with Rgl2^RA^ contrasts with the on/off kinetics for KRas4B^WT^ and KRas4B^G12V^ mutant with BRAF^RBD^, a well-established Ras effector. In that case, BRAF^RBD^ interacted with both the wildtype and the G12V mutant in a similar manner (Fig. 1C). The fitting using a 1:1 binding model was adequate for both KRas4B^WT^ and KRas4B^G12V^ cases (the RSS values are about 6.75 and 13.2, respectively, Fig. 1C) and the 2:1 binding model only marginally improved the fitting (the RSS values are about 2.96 and 4.34, respectively, Supplementary Fig. S2C).

These results indicate that the G12V oncogenic mutation may impact more significantly on the Rgl2-mediated signalling pathway and highlight the distinct binding mode of KRas4B^WT^ and KRas4B^G12V^ mutant to Rgl2^RA^ compared to BRAF^RBD^.

The circular dichroic (CD) spectra for KRas4B^WT^ and KRas4B^G12V^ samples across the increasing temperature from 20°C to 90°C showed improved structural stability for KRas4B^G12V^, which might contribute to the altered binding kinetics towards Rgl2^RA^ (Fig. 1 D-F).

### A 2:2 tetramer of KRas4B^G12V^:Rgl2^RA^ complex in the crystal structure

We conducted crystallization trials to obtain structural insights into the KRas4B:Rgl2^RA^ complex. To purify the complex, a mixture of GMPPNP-loaded KRas4B and Rgl2^RA^ was applied onto a size-exclusion column. The major elution peak corresponded to the co-elution of KRas4B^G12V^ and Rgl2^RA^ as a complex, and, as expected, the complex eluted earlier than KRas4B^G12V^ or Rgl2^RA^ alone (Supplementary Fig. S1C).

Crystals of the KRas4B^G12V^:Rgl2^RA^ complex were obtained which diffracted at a resolution of 3.07Å (Table 1). The phase was solved by molecular replacement using the structure of the human HRas^E31K^:rat RALGDS^RA^ complex (PDB ID 1LFD) (Huang *et al*., 1998). The space group was assigned to P1211 with four proteins per asymmetric unit (two molecules of KRas^G12V^ and two molecules of Rgl2^RA^) arranged as a tetramer (Fig. 2). The overall arrangement of KRas4B^G12V^:Rgl2^RA^ crystal structure is similar to HRas^E31K^:RALGDS^RA^ crystal structure (PDB ID 1LFD (Huang *et al*., 1998))(Supplementary Fig. S3). The complex forms a 2:2 heterotetramer where β2 (within Switch I) of KRas^G12V^ and β2 of Rgl2^RA^ generate a continuously extended β-sheet, along with interaction at Switch II of the same KRas^G12V^ molecule with the second Rgl2^RA^ molecule (Fig. 3 A, B and C and Fig. 4 A and B). In both KRas4B^G12V^:Rgl2^RA^ and HRas^E31K^:RALGDS^RA^ cases, the Ras-Ras interface is formed within the region spanning amino acids 1-90 where KRas4B^G12V^ and HRas^E31K^ share 100% amino acid sequence identity except for the point mutations G12V and E31K. The KRas4B^G12V^:Rgl2^RA^ heterotetramer complex is stabilized by a network of hydrogen bonds and hydrophobic interactions (summarized in Supplementary Fig. S4A), leading to complimentary surface charges (Supplementary Fig. S4B).

**Table 1.** Data collection and refinement statistics for human KRas4B^G12V^:Rgl2^RA^ complex. Statistics for the highest-resolution shell are shown in parentheses.

|  |  |
| --- | --- |
| Wavelength | 0.9795 |
| Resolution range | 28.68 - 3.071 (3.18 - 3.071) |
| Space group | P 32 2 1 |
| Unit cell | 77.843 77.843 163.728 90 90 120 |
| Total reflections | 106507 (7598) |
| Unique reflections | 11107 (949) |
| Multiplicity | 9.6 (8.0) |
| Completeness (%) | 98.22 (86.51) |
| Mean I/sigma(I) | 11.15 (1.64) |
| Wilson B-factor | 95.59 |
| R-merge | 0.1313 (0.6732) |
| R-meas | 0.1389 (0.7159) |
| R-pim | 0.04474 (0.2379) |
| CC1/2 | 0.998 (0.957) |
| CC* | 1 (0.989) |
| Reflections used in refinement | 11065 (936) |
| Reflections used for R-free | 582 (41) |
| R-work | 0.2367 (0.3229) |
| R-free | 0.3204 (0.3572) |
| CC(work) | 0.943 (0.897) |
| CC(free) | 0.851 (0.486) |
| Number of non-hydrogen atoms | 4116 |
| macromolecules | 4050 |
| ligands | 66 |
| solvent | 0 |
| Protein residues | 514 |
| RMS(bonds) | 0.007 |
| RMS(angles) | 1.40 |
| Ramachandran favored (%) | 95.22 |
| Ramachandran allowed (%) | 3.98 |
| Ramachandran outliers (%) | 0.80 |
| Rotamer outliers (%) | 0.89 |
| Clashscore | 14.47 |
| Average B-factor | 113.50 |
| macromolecules | 113.74 |
| ligands | 98.55 |

**Figure 3.**
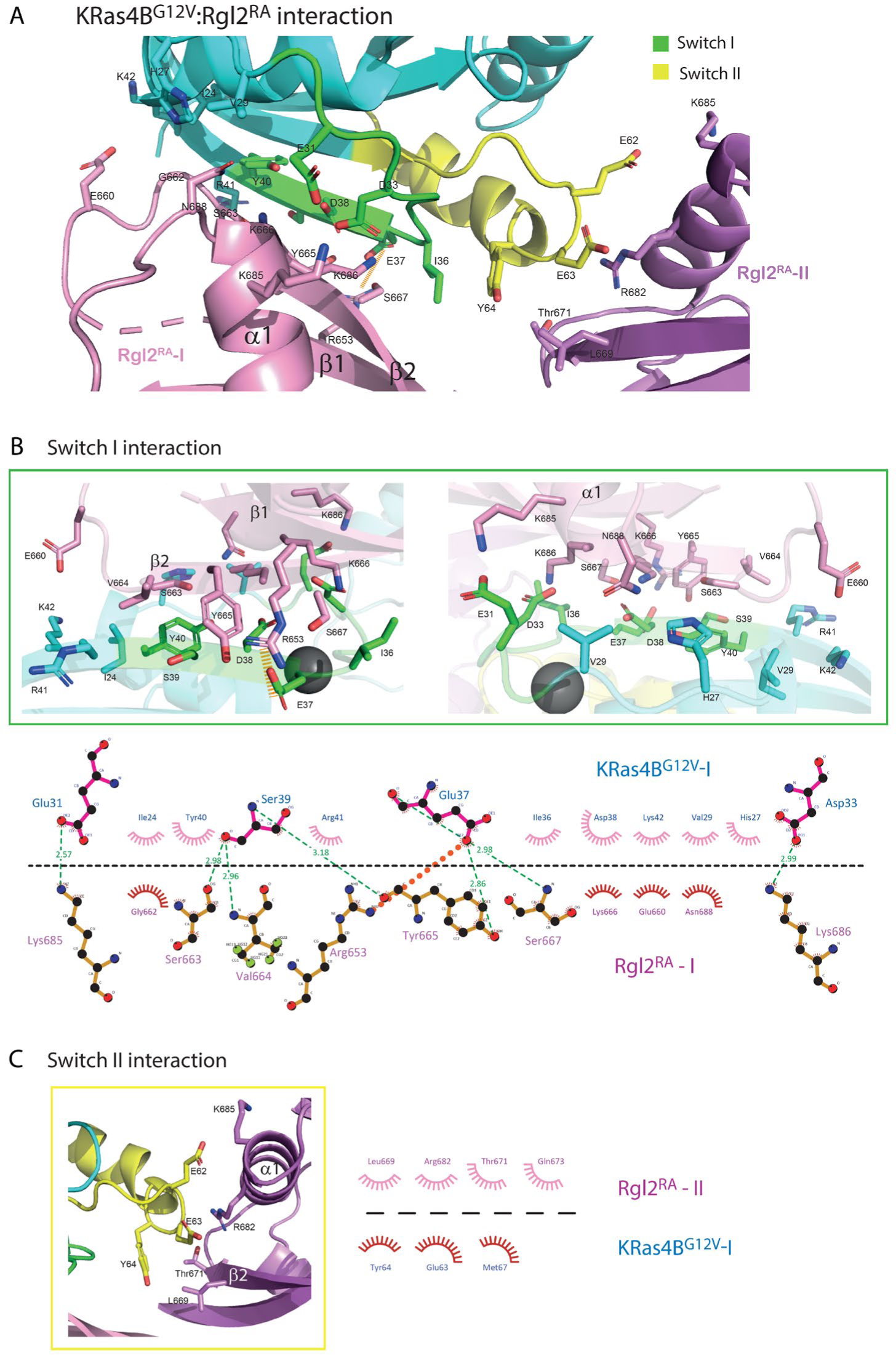
The interacting interface of KRas4B^G12V^ and Rgl2^RA^ of the KRas4B^G12V^:Rgl2^RA^ 2:2 heterotetramer. (A) The overview of the interacting interface of KRas4B^G12V^ and Rgl2^RA^, highlighting the residues involved in hydrogen and hydrophobic interactions between β1, β2 and α1 of Rgl2^RA1^ (violet sticks) and KRas4B^G12V^ Switch-I (green sticks, enlarged in the green box) or Switch-II (yellow sticks, enlarged in the yellow box). An orange dashed line shows a salt bridge formed between E37 of KRas4B^G12V^ and R653 of Rgl2^RA^. (B) Upper panel: Blow-up images of the KRas4B^G12V^:Rgl2^RA^ interface involving Switch-I. Lower panel: a schematic representation of the intermolecular contacts where hydrogen bonds (green dashed lines), a salt bridge (orange dashed line) and hydrophobic contacts (spiked arches) were predicted by LIGPLOT (Wallace *et al*., 1995) (lower panel). Numbers indicate atomic distances in Å. (C) A blow-up image of the KRas4B^G12V^:Rgl2^RA^ interface involving Switch II (left panel) and its LIGPLOT representation (right panel).

**Figure 4.**
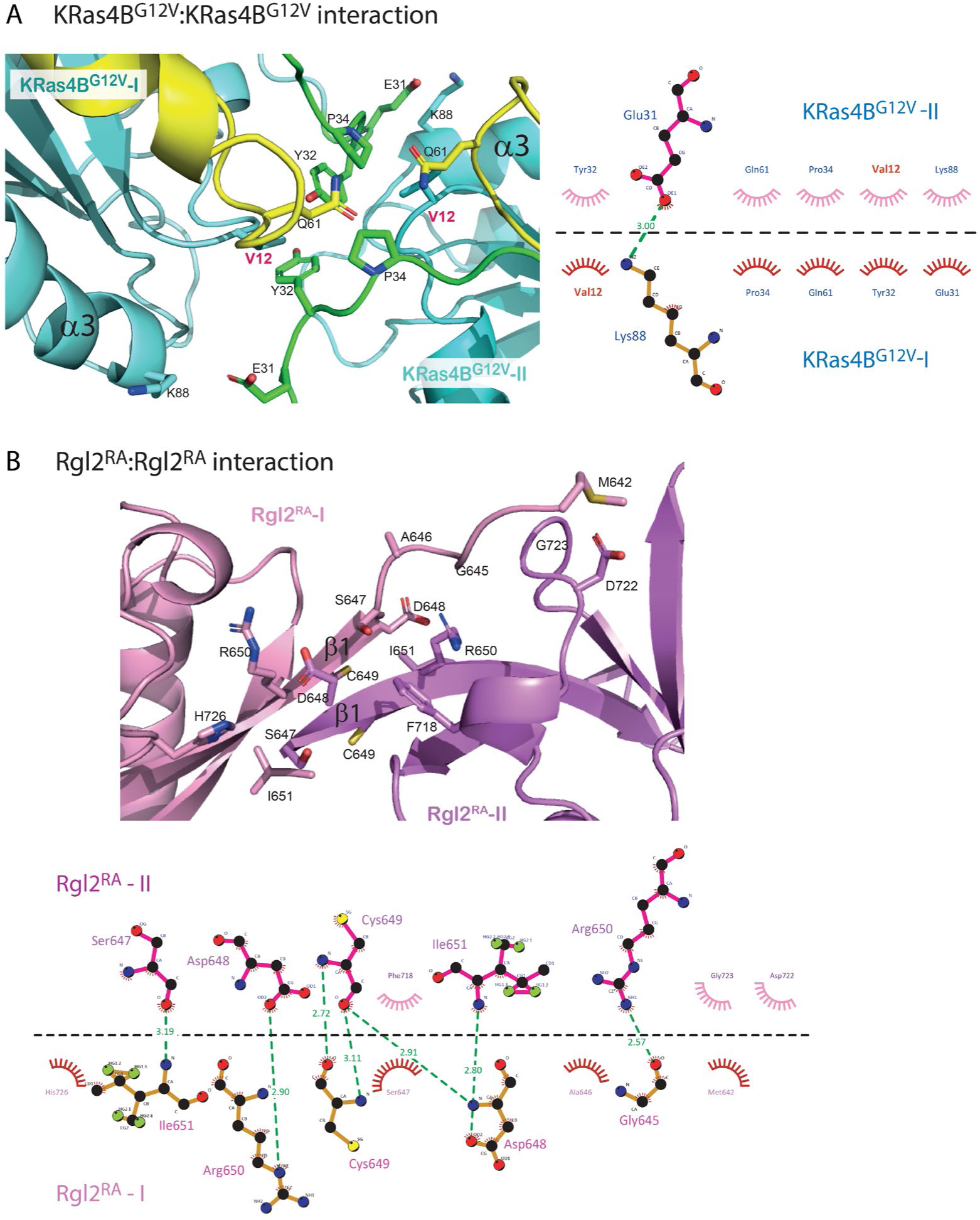
The interacting interface of KRas4B^G12V^: KRas4B^G12V^ and Rgl2^RA^: Rgl2^RA^ of the KRas4B^G12V^:Rgl2^RA^ 2:2 heterotetramer. (A) The interface of KRas4B^G12V^: KRas4B^G12V^, highlighting the residues involved in hydrogen and hydrophobic interactions between two KRas4B^G12V^ molecules. The interface comprises Switch-I, Switch-II and α3 of the two KRas4B^G12V^ molecules. The oncogenic mutation, V12, is annotated in red letters. A schematic representation of the intermolecular contacts predicted by LIGPLOT is presented in the right panel. (B) The interacting interface of Rgl2^RA^: Rgl2^RA^, highlighting the residues involved in hydrogen and hydrophobic interactions between two Rgl2^RA^ molecules, shaded in pink and violet. The interaction occurs at the N-terminal between the two anti-parallel β1. A LIGPLOT diagram is shown below.

### KRas^G12V^:Rgl2^RA^ interface

KRas4B^G12V^ residues within the Switch I interact with residues in β1, β2 and α1 of Rgl2^RA^. These interactions include a salt bridge between E37 of KRas4B^G12V^ Switch I and R653 of Rgl2^RA^ (Fig. 3A and B). The Switch II of the same KRas4B^G12V^ molecule also contributes to the complex formation by interacting with the second Rgl2^RA^ through residues in β2 and α1 (Fig. 3A and C).

### KRas^G12V^: KRas^G12V^ interface

Meanwhile, two KRas4B^G12V^ molecules have direct contact through Switch I, Switch II and α3 (Fig. 4A, and Supplementary Fig. S4A). Importantly, the V12 residue of the oncogenic G12V mutation is in the proximity of the ring of Y32 of the neighbouring KRas4B^G12V^, contributing to the KRas4B^G12V^: KRas4B^G12V^ interface, increasing the hydrophobic pocket.

### Rgl2^RA^:Rgl2^RA^ interface

The β1 of both Rgl2^RA^ molecules run anti-parallel to each other, interacting through various hydrophobic interactions and hydrogen bonds at both side-chain and backbone levels (Fig. 4B, and Supplementary Fig. S4A).

### Solution NMR analyses of the KRas4B^G12V^:Rgl2^RA^ complex

The KRas4B^G12V^:Rgl2^RA^ complex was further analyzed in solution by nuclear magnetic resonance (NMR). First, the solution structure of free Rgl2^RA^ was determined by solution NMR (Supplementary Fig. S5A) (Table 2). As expected, the structure adopts the ββαββαβ ubiquitin-fold structure, a common feature for the RA/RBDs (Kiel and Serrano, 2006). Overall, it is similar to the structure of Rlf, the mouse homologue of human Rgl2 (PDB ID 1RLF)(Esser *et al*., 1998) (Supplementary Fig. S5B). The solution structure of free Rgl2^RA^ is also very similar to the crystal structure of Rgl2^RA^ in the KRas4B^G12V^:Rgl2^RA^, indicating that the complex formation causes relatively small structural changes, and the crystal structure likely reflects the physiological Rgl2^RA^ folding (Fig. 5A).

**Table 2.** NMR structure statistics for Rgl2^a^.

| Quantity |  |
| --- | --- |
| Assigned <sup>1</sup>H/<sup>13</sup>C/<sup>15</sup>N chemical shifts | 305/256/56 |
| NOE distance restraints<sup>b</sup> | 866/281/777 |
| Max. distance restraint violation (Å) | 0.13 ± 0.01 |
| Deviations from idealised geometry: |  |
| Bond lengths (Å) | 0.0138 ± 0.0001 |
| Bond angles (°) | 1.81 ± 0.03 |
| AMBER energy (kcal/mol) | −3624.25 ± 78.09 |
| AMBER vdW energy (kcal/mol) | −279.71 ± 12.49 |
| Ramachandran plot statistics<sup>c</sup> (%) | 89.5/10.5/0.1/0 |
| Backbone RMSD (Å)<sup>d</sup> | 0.42 ± 0.05 |
| All heavy atom RMSD (Å)<sup>d</sup> | 0.84 ± 0.06 |
| Backbone RMSD to the reference (Å)<sup>e</sup> | 0.91 |
| All heavy atom RMSD to the reference (Å)<sup>e</sup> | 1.68 |

<sup>a</sup>Where applicable, the average value and the standard deviation over the 20 energy-refined conformers obtained by the program OPALp 2.1 in the presence of the experimental restraints. CYANA calculations were started from 100 conformers with random torsion angle values, simulated annealing with 10,000 torsion angle dynamics steps was applied.
<sup>b</sup>Short/medium/long-range distance restraints derived from NOESY spectra.
<sup>c</sup>Percentage of residues in the most favoured/additionally allowed/generously allowed/disallowed regions of the Ramachandran plot according to the program PROCHECK.
<sup>d</sup>RMSD to the mean structure for residues 649–657, and 665–732.
<sup>e</sup>RMSD between the closest structure to the reference among the ensemble conformations and the reference structure (the crystal structure of the KRas4B<sup>G12V</sup>:Rgl2 complex).

**Figure 5.**
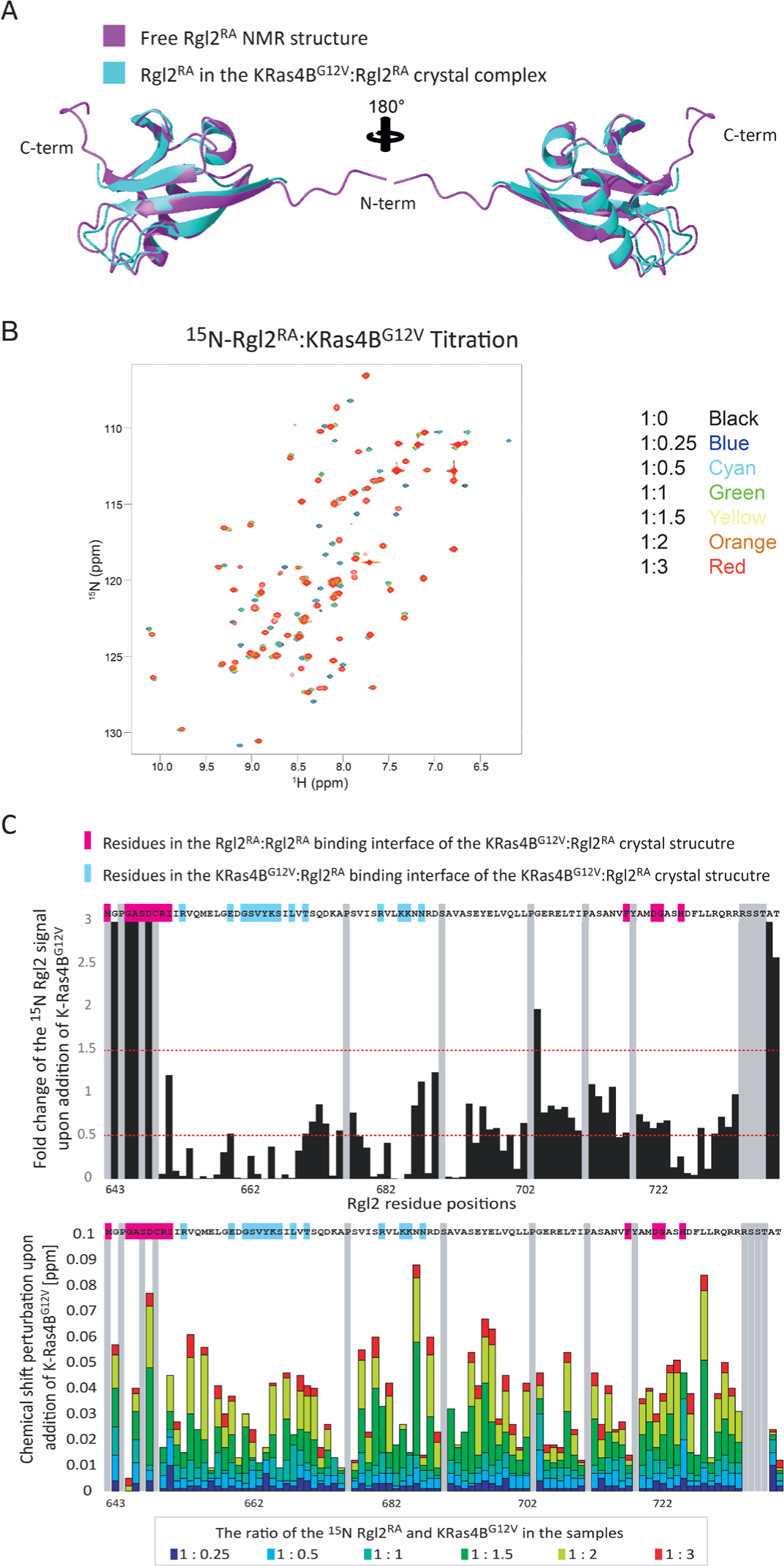
NMR analysis of KRas4B^G12V^:Rgl2^RA^ complex formation in solution. (A) Comparison between the Rgl2^RA^ structures identified in solution NMR (shown in magenta) and in crystal complex with KRas4B^G12V^ (shown in cyan). (B)-(D) ^1^H-^15^N HSQC titration analysis of ^15^N-labelled Rgl2^RA^ upon addition of non-labelled KRas4B^G12V^ supports the KRas4B^G12V^:Rgl2^RA^ tetramer formation. (B) Overlays of 2D ^1^H-^15^N HSQC NMR spectra from multipoint titrations of ^15^N-labelled Rgl2^RA^ with non-labelled KRas4B^G12V^. Rgl2^RA^:KRas4B^G12V^ molar ratios of the titration samples are colour-coded as follows; 1:0 - black, 1:0.25 - blue, 1:0.25 – cyan, 1:0.5 – green, 1:1 – yellow, 1:2 – orange and 1:3 – red. (C) The NMR signal intensity changes (upper panel) and the chemical shift perturbation (lower panel) of backbone ^1^H^N^ and ^15^N nuclei of Rgl2^RA^ with non-labelled KRas4B^G12V^, presented in (B), are summarised as column diagrams as a function of Rgl2^RA^ amino acid sequence. Proline and unassigned Rgl2^RA^ residues are shaded in grey. Rgl2^RA^ residues involved in the Rgl2^RA^:Rgl2^RA^ interface of the KRas4B^G12V^:Rgl2^RA^ crystal structure are highlighted in pink on the amino acid sequence, and residues involved in the KRas4B^G12V^:Rgl2^RA^ interface are highlighted in blue. Rgl2^RA^ residue position numbers, according to the UniProt, are indicated at the bottom of the diagram. (Upper panel) The signal intensities of Rgl2^RA^ residues in the presence of three times molar excess of KRas4B^G12V^ were divided by the signal intensities in the absence of KRas4B^G12V^ and plotted as a bar-chart graph. Red-dotted lines are drawn at the fold-change values of 0.5 and 1.5 to highlight the residues that show a substantial increase or decrease of the signals upon the addition of KRas4B^G12V^. (Lower panel) The chemical shift perturbation of backbone ^1^H^N^ and ^15^N nuclei of Rgl2^RA^ with non-labelled KRas4B^G12V^. The mean shift difference Δδave was calculated as [(Δδ^1^H^N^)^2^ + (Δδ^15^N/10)^2^]^1/2^ where Δδ^1^H^N^ and Δδ^15^N are the chemical shift differences between Rgl2^RA^ on its own and in the presence of non-labelled KRas4B^G12V^. The bar graphs are colour-coded according to the Rgl2^RA^- KRas4B^G12V^ concentration ratio.

The KRas4B^G12V^:Rgl2^RA^ complex was next analyzed in solution by NMR chemical shift perturbation. Two-dimensional (2D) ^15^N–^1^H-heteronuclear single quantum coherence (HSQC) spectra were measured for the ^15^N-labelled Rgl2^RA^ sample in the absence or presence of an increasing amount of non-labelled KRas4B^G12V^ (Fig. 5B and C). Chemical shift perturbations were observed for many of the Rgl2^RA^ residues, in agreement with KRas4B^G12V^:Rgl2^RA^ complex formation in solution. Most of the Rgl2^RA^ residues at the KRas4B^G12V^:Rgl2^RA^ and Rgl2^RA^:Rgl2^RA^ interfaces in the crystal structure showed greater changes (either decreased or increased) in their NMR signal intensities and in chemical shift perturbation, indicating their participation in the complex formation and supporting the heterodimer interface observed in the crystal structure (Fig. 5B and C). Signals from the Rgl2^RA^ residues in β1 at the N-terminal end (642-649) display the largest changes. This is in agreement with the KRas4B^G12V^:Rgl2^RA^ heterotetramer crystal structure where the highly flexible N-terminal region of free Rgl2^RA^ (Supplementary Fig. S5A) becomes rigid through interaction with another Rgl2 molecule in the heterotetramer (Fig. 4B). In addition, the overall decrease in the NMR signals is compatible with the formation of a relatively large complex such as the KRas4B^G12V^:Rgl2^RA^ heterotetramer complex (62kDa). Meanwhile, the titration experiment showed that the chemical shift changes were not fully saturated in most of the Rgl2^RA^ residues when the Rgl2^RA^:Kras^G12V^ molar ratio was 1:1 or even 1:2 (Fig. 5C), suggesting that the stoichiometry is more complex than simple 1:1, possibly due to the heterotetramer formation.

We next examined the NMR chemical shift perturbation of KRas4B^G12V^ in solution in the absence or presence of non-labelled Rgl2^RA^ (Figure 6). When loaded with GMPPNP, the number of detectable peaks of the ^15^N labelled KRas4B^G12V^ signal is substantially decreased compared to the number of signals obtained from the GDP-loaded sample (Supplementary Fig. S6) as previous studies reported (Ito et al., 1997; Menyhard et al., 2020). Consequently, unfortunately, most of the KRas4B^G12V^ residues at the KRas4B^G12V^:Rgl2^RA^ complex interface could not be detected (Fig. 6A and B), except for K42 and K88, both of which showed substantially decreased signal intensities upon Rgl2^RA^ addition, in agreement with their expected participation in the complex formation (Fig. 6C). Furthermore, the addition of Rgl2^RA^ leads to a decrease of most of the KRas4B^G12V^ signals further suggesting formation of a relatively large complex, like the heterotetramer.

**Figure 6.**
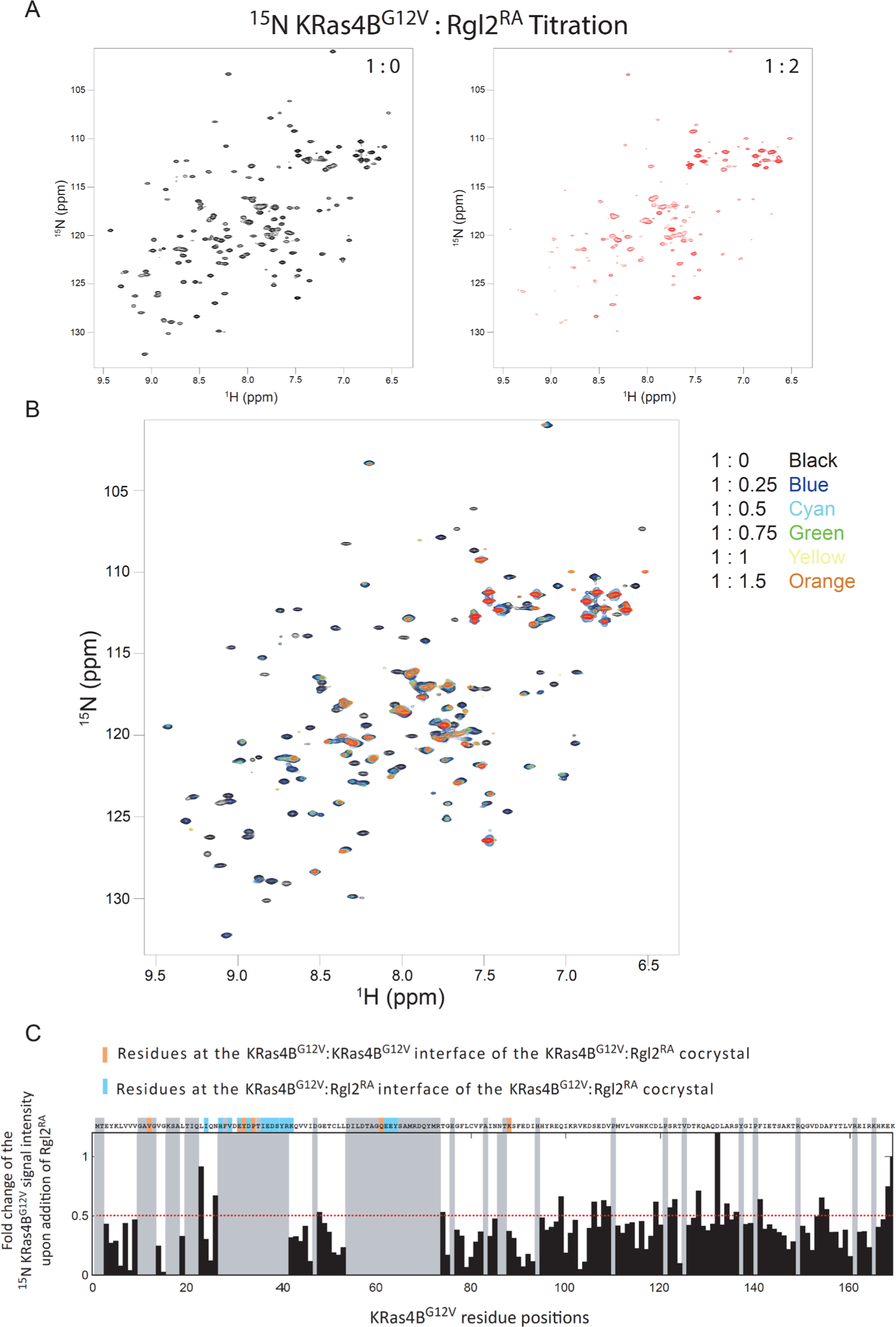
KRas4B^G12V^:Rgl2^RA^ complex formation in solution. ^1^H-^15^N-HSQC titration analysis of ^15^N-labelled KRas4B^G12V^ upon addition of non-labelled Rgl2^RA^ supports the KRas4B^G12V^:Rgl2^RA^ tetramer formation. (A) and (B) The 2D ^1^H-^15^N-HSQC NMR spectra of KRas4B^G12V^. ^15^N-labelled KRas4B^G12V^ was titrated with non-labelled Rgl2^RA^. (A) The 2D ^1^H-^15^N-HSQC spectra of KRas4B^G12V^: Rgl2^RA^ complex mixed with the molar ratio of 1:0 (black, left panel) and 1:2 (red, right panel) are shown. Many signals from ^15^KRas4B^G12V^ residues disappeared upon the addition of Rgl2^RA^. (B) Superimposed 2D ^1^H-^15^N-HSQC NMR spectra of ^15^N- labelled KRas4B^G12V^:Rgl2^RA^ titration experiments. The titration samples are colour-coded as follows; 1:0-black, 1:0.25 - blue, 1:0. 5 – cyan, 1:0.75 –green, 1:1 – yellow, 1:1.5 – orange and 1:2 – red. (C) Fold changes of the signal intensities of ^15^N-labelled KRas4B^G12V^ upon the addition of non-labelled Rgl2^RA^. The signal intensities of KRas4B^G12V^ residues in the presence of two times molar excess of Rgl2^RA^ were divided by the signal intensities in the absence of Rgl2^RA^, and the obtained values were plotted as a column graph. Undetectable residues are shaded in grey. The residue T2 was also shaded grey, as the chemical shift after the addition of Rgl2^RA^ overlapped with other signals. A red-dotted line is drawn at the fold-change values of 0.5 to indicate that most residues show a substantial decrease in the signals upon the addition of Rgl2^RA^. KRas4B^G12V^ residues in the KRas4B^G12V^: KRas4B^G12V^ interface of the KRas4B^G12V^:Rgl2^RA^ crystal structure is highlighted in orange, and residues in the KRas4B^G12V^:Rgl2^RA^ interface of the crystal structure is highlighted in blue. KRas4B^G12V^ residue positions according to the UniProt are indicated at the bottom of the diagram.

To estimate the size of the complex, we conducted NMR relaxation measurements. The ^15^N longitudinal (*T*1) and transverse (*T*2) relaxation times, and the steady-state heteronuclear {^1^H}- ^15^N NOE of ^2^H/^13^C/^15^N- labelled KRas4B^G12V^, mixed with non-labelled Rgl2^RA^, was measured with TROSY-based pulse schemes (Supplementary Fig. S7A)(Lakomek et al., 2012). The overall rotational correlation time *τ*c deduced from the measured *T1*, *T2* and NOE was approximately 14.7 nsec. Meanwhile, the *τ*c values estimated theoretically for the dimer and the tetramer were approximately 14.2 nsec and 38.8 nsec when we assumed that they were spherical and the radii of gyration were 25 Å and 35 Å, respectively (Supplementary Fig. S7B). These results suggest that the status of the KRas4B^G12V^-Rgl2^RA^ complex in solution may be closer to the heterodimer structure. However, since the crystal structures are ellipsoidal rather than spherical, and an equilibrium-like exchange process between the heterodimer and heterotetramer, or transient tetramer formation, may occur, further NMR relaxation measurements and detailed analysis would be needed to more accurately predict the complex status.

### Mass photometry indicates the presence of a heterotetramer of KRas4B^G12V^:Rgl2^RA^ complex

To address the question of whether KRas4B^G12V^:Rgl2^RA^ complex can exist as a heterotetramer in solution, we analyzed the complex by mass photometry, a label-free technique that has recently been adapted to measure the mass of biomolecules in solution (Young et al., 2018). Our commercially-available instrument (see Materials and methods) can measure masses in the range 30-5000 kDa, and is suitable to detect KRas4B^G12V^:Rgl2^RA^ heterotetramer (62kDa). The KRas4B^G12V^:Rgl2^RA^ complex was freshly prepared by size exclusion chromatography at 4°C, and the peak fraction that contained both proteins at 1:1 ratio was used for the measurement (Fig. 7A). The histogram of the frequency counts for this sample showed a peak corresponding to a biomolecular complex with a molecular weight of 68 kDa (Fig. 7B). As the expected molecular weight of the KRas4B^G12V^:Rgl2^RA^ heterotetramer is about 62 KDa, the result indicates that at least part of the complex population likely exists as the 2:2 heterotetramer in solution.

**Figure 7.**
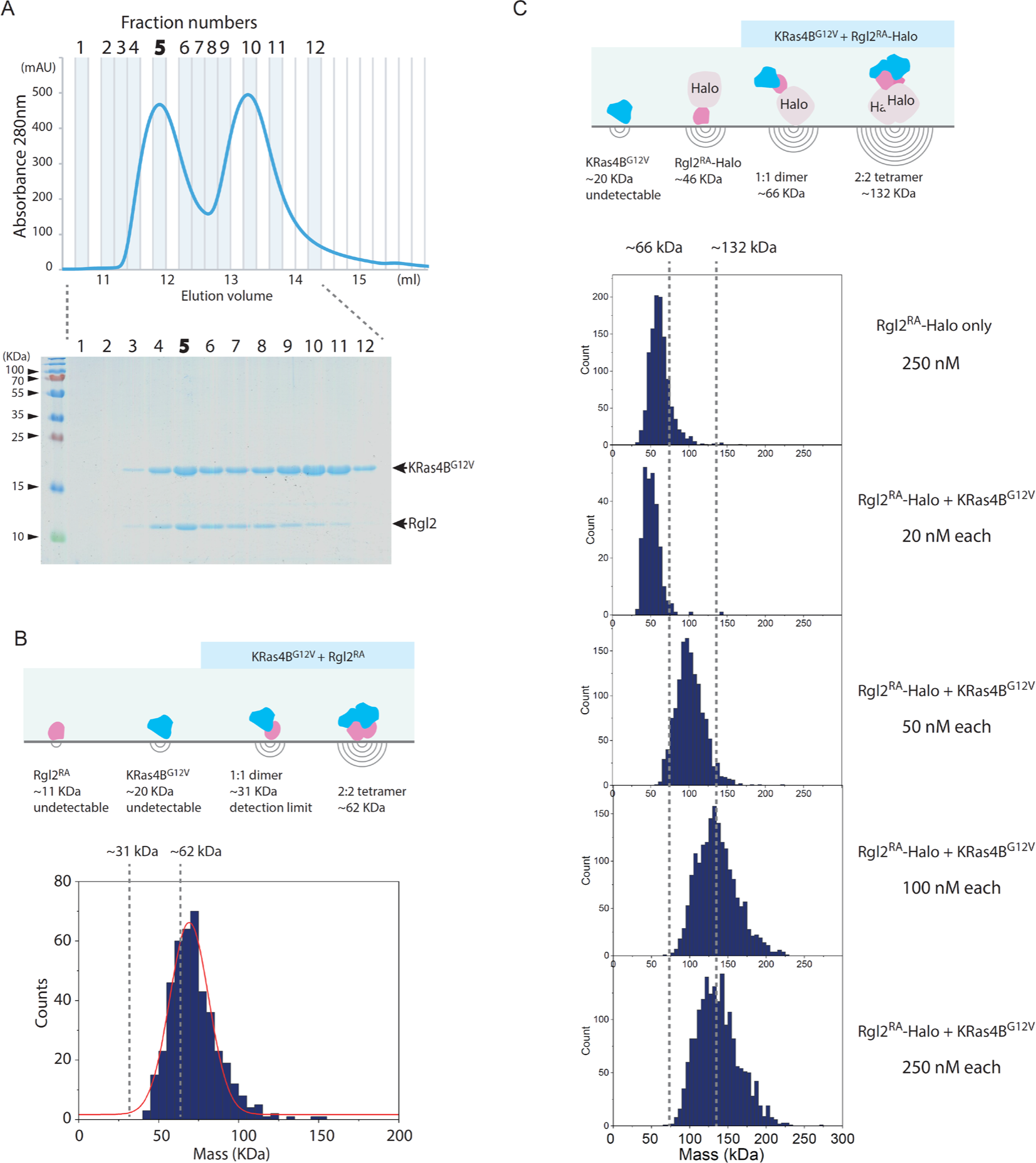
KRas4B^G12V^:Rgl2^RA^ complex can form a heterotetramer in solution. Mass photometry analysis of KRas4B^G12V^:Rgl2^RA^ complex in solution. (A) KRas4B^G12V^:Rgl2^RA^ complex was purified using size exclusion chromatography. The fractions were analysed with 15% SDS-PAGE gel (upper panel) according to the elution profile (lower panel). Fraction 5 was chosen and measured using mass photometry (OneMP, Refeyn). (B) The cartoon representing the different complex configurations possibilities indicates that only heterodimers (∼31kDa) and heterotetramers (∼62kDa) can be detected by the system. Histogram of the frequency counts against the purified KRas4B^GV^:Rgl2^RA^ complex with fitting for Gaussian distribution (red). The duration of the video analysed was 60s, and it shows that the population identified has an average mass of ∼68kDa, which is in good agreement with the expected MW of the heterotetramer (61.7kDa). (C) KRas4B^G12V^ and Halo-tagged Rgl2^RA^ (Rgl2^RA^-Halo) complex formation. The cartoon represents MWs of possible complexes. KRas4B^G12V^ and Rgl2^RA^-Halo were separately prepared and were mixed to generate a premix 2 µM sample, which was further diluted to 20 nM, 50 nM, 100 nM and 250 nM before the measurement. At 250 nM, Rgl2^RA^-Halo without KRas4B^G12V^ showed about 55 kDa, within the range of the predicted MW (∼46 kDa). In contrast, the mixed samples showed an increase in the MW as the concentration was increased. At 50nM, the observed MW was about 90 kDa, which may represent the predicted dimer, and at 100-250 nM, the observed MW peaked at about 130 kDa, which coincided with the predicted MW for the tetramer.

To detect the presence of the heterodimer, as well as the heterotetramer, we examined the complex formation between KRas4B^G12V^ and a Halo-tagged Rgl2^RA^ (Rgl2^RA^-Halo), where the Halo-tag increased the MW by 35kDa, hence the heterodimer of KRas4B^G12V^ and Rgl2^RA^-Halo (expected to be about 66 kDa) became within the detection range of the mass photometry instrument. The KRas4B^G12V^ and Rgl2^RA^-Halo were separately prepared and mixed at a 1:1 molar ratio at 2µM, and the 2µM premix sample was further diluted to 20, 50, 100 and 250 nM for the measurement (Fig. 7C). At 20 nM, the MW was about 50 kDa, in a similar range to the Rgl2^RA^- Halo monomer. At 50 nM, the MW increased to about 90 kDa, which may represent the dimer formation. At 100-250 nM, the peak of the MW reached about 130 kDa, which corresponded to the 2:2 hetero-tetramer. The result indicated that, under this experimental condition, the KRas4B^G12V^ and Rgl2^RA^-Halo may form a heterotetramer.

## Discussion

We conducted interaction studies of human KRas4B and Rgl2^RA^ to obtain structural insights into the oncogenic Ras-dependent activation of the RalA/B pathway. Our BLI data show that the oncogenic mutation G12V alters the binding kinetics between KRas4B and Rgl2^RA^. Using X-ray crystallography, we found that KRas4B^G12V^ and Rgl2^RA^ form a heterotetramer and that the oncogenic G12V mutation resides at the interaction surface. Meanwhile, NMR and mass photometry analyses indicate the complex exists in a heterogeneous status in solution.

### The KRas4B^G12V^:Rgl2^RA^ dimer formed through the KRas4B^G12V^ Switch I region displays the common features observed in the canonical mode of Ras-effector interaction

The KRas4B^G12V^:Rgl2^RA^ contacts involving the KRas4B^G12V^ Switch I region share the common structural feature as the canonical Ras-effector interface; the β2 in the Switch I region of Ras and β2 of RBD/RA interact to form a parallel β-sheet (Supplementary Fig. S8). However, the KRas4B^G12V^ and Rgl2^RA^ molecules at this contact are situated at a slightly different angle compared to Kras:RBD complexes, such as KRas4B^WT^:CRAF^RBD^ (PDB ID 6VJJ) (Supplementary Fig. S8). This is consistent with the recent structural evaluation of all available RAS:RA/RBD complex structures (Eves et al., 2022). In this respect, KRas4B^G12V^:Rgl2^RA^ complex belongs to the “RalGDS- family cluster”, involving HRas^E31K^:RALGDS^RA^ (PDB:1LFD) and KRas^WT/G12V^:Rgl1^RA^ (PDB: 7SCW and 7SCX) complexes (Eves *et al*., 2022), rather than to the “CRAF cluster”, suggesting a structure-function correlation.

### The interaction mode between Ras and Rgl2/RALGDS is distinct from other Ras:RBD/RA interactions

The key contributor of the heterotetramer formation seen in our structure of KRas4B^G12V^:Rgl2^RA^, and the previously published HRas^E31K^:RALGDS^RA^ is the interaction between the Switch II region of Ras and the second RBD/RA molecule (Fig. 3C, and Supplementary Fig. S9A). This feature is distinct from other previously reported Ras:effector crystal structures, where the Switch II region is either not participating in the interaction (Ras:CRAF^RBD^, PDB ID 6VJJ(Tran et al., 2021)) or is interacting with the same RBD that interacts with the Switch I region of the same Ras molecule (Ras:PI3Kγ^RBD^ (PDB ID 1HE8)(Pacold et al., 2000), Ras: PLCε^RA^ (PDB ID 2C5L)(Bunney et al., 2006), Ras:RASSf5^RA^ (PDB ID 3DDC)(Stieglitz et al., 2008)), hence forming a Ras:RA/RBD heterodimer, rather than a heterotetramer (Supplementary Fig. S9B).

One interesting case is a recently reported structural study of Rgl1, yet another RalGEF (Eves *et al*., 2022). The primary sequences of Rgl1^RA^, Rgl2^RA^ and RALGDS^RA^ are distinct but close to each other when compared to CRAF^RBD^, a representative RBD (Supplementary Fig. S10). The crystal structures of KRas4B^WT^ or KRas4B^G12V^ in complex with Rgl1^RA^ show that Rgl1^RA^ interact with the Switch I region of Ras in a highly similar manner as other complexes; the similarity is particularly high when compared to K-Ras4B^G12V^:Rgl2^RA^ (this study) and HRas^E31K^:RALGDS^RA^ (PDB ID 1LFD) (Supplementary Fig. S9B and Fig. S11 top row). In addition, Rgl1^RA^ contact the Switch II region in a manner similar to PI3Kγ^RBD^ and PLCε^RA^ (Supplementary Fig. S9B). However, this makes Rgl1^RA^ distinct from other RasGEFs, Rgl2^RA^ and RalGDS^RA^, as these establish the Switch II contact using the second RA molecule (Supplementary Fig. S9A and Fig. S11).

The crystal structure of Rgl1 in complex with KRas4B^G12V^ (PDB ID 7SCX) also suggests a possible heterotetramer formation (Eves *et al*., 2022). However, the proposed structural arrangements of the KRas4B^G12V^:Rgl1^RA^ heterotetramer is distinct from the Ras:Rgl2/RALGDS heterotetramers (supplementary Fig. S11); in the Ras4B^G12V^:Rgl1^RA^ structure, the heterodimer is mediated by an additional Rgl1^RA^:Rgl1^RA^ interaction but the two KRas^G12V^ molecules are not in contact with each other. Indeed, the second Rgl1^RA^ associates with the first Rgl1^RA^ through its C-terminal end, which extends into the paired Rgl1^RA^ in a reciprocal manner, causing a “domain swap” of the β5 (Eves *et al*., 2022)(Supplementary Fig. S11). Consequently, the second KRas^G12V^ in the KRas^G12V^:Rgl1^RA^ heterotetramer is away from the first KRas^G12V^ molecule, and the G12V mutation is not directly involved in the molecular contacts. In agreement with this structural arrangement, the affinity between KRas and Rgl1^RA^ was reported unaffected by the G12V oncogenic mutation (Eves *et al*., 2022). This is distinct from the heterotetramer observed in the case of Rgl2^RA^, where V12 is directly involved in the RAS:RAS interaction, in agreement with the altered binding kinetics of KRas^G12V^ to Rgl2^RA^ in comparison to KRas^WT^ (Fig. 4A, Supplementary Fig. S4 and Supplementary Fig. S11).

### Oligomerisation of KRas:effector complexes

Accumulating evidence suggests that KRas is capable of forming a dimer in solution even in the absence of effectors (Andreadelis et al., 2022; Ingolfsson et al., 2022; Jang et al., 2016; Lee et al., 2021; Lee et al., 2020; Muratcioglu et al., 2015; Ozdemir et al., 2022; Packer et al., 2021; Prakash et al., 2017; Sarkar-Banerjee et al., 2017). The physiological importance of Ras dimerisation and its potential to be a therapeutic target was highlighted by a G12D-specific inhibitor, BI2852, which may cause artificial dimerisation and block the protein function (Kessler et al., 2019; Tran et al., 2020). Furthermore, Ras forms nanoclusters in the membrane *in vivo* and *in vitro* (Lakshman et al., 2019; Plowman et al., 2005; Prior et al., 2003; Weise et al., 2011; Zhou et al., 2014). The modes of these Ras oligomer formations are dependent on various parameters, including membrane lipid compositions, Ras nucleotide binding status, availability of Ras effectors and the actin cytoskeleton and are expected to be context-dependent.

Previously it has been suggested that Ras in general can form dimers in solution but structural information on these dimers vary and the dimerisation status may therefore be categorised into four classes based on the α helices and β sheets at the interface as follows; (i) α4/α5 (Andreadelis *et al*., 2022; Jang *et al*., 2016; Lee *et al*., 2021; Lee *et al*., 2020; Packer *et al*., 2021; Prakash *et al*., 2017), (ii) α3/α4 (Jang *et al*., 2016; Muratcioglu *et al*., 2015; Prakash *et al*., 2017), (iii) α/β (Lee *et al*., 2021), and (iv) β/β (Jang *et al*., 2016; Muratcioglu *et al*., 2015) (Supplementary Fig. S12). Notably, the KRas4B^G12V^:KRas4B^G12V^ interface that we observe in the KRas4B^G12V^:Rgl2^RA^ heterotetramer does not belong to any of these categories and uses the unstructured regions of Switch I, Switch II, V12 in the P-loop and K88 (Fig. 4A, and Supplementary Fig. S12), in a similar manner to the HRas^E31K^:RALGDS^RA^ crystal (PDB ID 1LFD) (Huang *et al*., 1998). While this mode of tetramer formation could be an outcome of a crystal-packing artefact (Vetter et al., 1999), it is worth noting that the spatial arrangement of symmetry mates of human HRas^WT^ crystal (PDB ID 5P21) also shows a similar Ras:Ras contact (Pai et al., 1990) (Supplementary Fig. S13C). Therefore, this Ras:Ras contact mode may reflect the intrinsic nature of Ras molecules, which may be stabilized by both KRas4B 12V and Rgl2^RA^ to form the heterotetramer.

Our mass photometry analysis supports a tetramer formation in solution. However, the rotational correlation time from the NMR relaxation experiments was rather close to that estimated from the size of the heterodimer. Furthermore, our SEC-MALS attempt only detected a heterodimer formation (data not shown), indicating that the heterotetramer formation may involve certain environmental conditions or an exchange process, as was the case for the KRas^GV^:Rgl1^RA^ where tetramerization was observed by SEC only on 6-months old samples (Eves *et al*., 2022). Physiological settings, including the presence of the plasma membrane and the full-length effector molecules rather than just the RBD/RA, may stabilize higher-order complex formations.

### Possible involvement of V12 in the KRas4B:Rgl2^RA^ complex formation and oncogenicity

The KRas4B^G12V^:KRas4B^G12V^ interface of the KRas4B^G12V^:Rgl2^RA^ tetramer complex involves the residue V12, an oncogenic amino-acid substitution occurring in about 28% of mutated *KRAS* cases (COSMIC). This G12V substitution alters the binding kinetics towards Rgl2^RA^ compared to the wildtype (Fig. 1B). In the KRas4B^G12V^:Rgl2^RA^ tetramer complex, the presence of V12 creates a larger hydrophobic pocket together with Y32, compared to the wildtype case of G12, seen in the HRas^WT^ crystal (PDB ID 5P21) or in the HRas^E31K^:RALGDS^RA^ crystal (PDB ID 1LFD) (Supplementary Fig. S13). Interestingly, the E31K substitution, used to stabilize the complex in the HRas^E31K^:RALGDS crystal structure (Huang *et al*., 1998) (Supplementary Fig. S13B), has been reported in cancer samples with *HRAS* mutations (COSMIC). Therefore, the capability of Ras mutants to form a stable complex with RalGEFs may be directly linked with Ras oncogenicity. It is interesting to note that in the previously proposed Ras:CRAF heterotetramer complex derived from SAXS data (Packer *et al*., 2021), the position of residue 12 is relatively distant from the interacting surfaces. An interesting speculation can be that the oncogenic substitution mutations at glycine 12 might have a greater impact on RalGEF-mediated signalling than Raf kinase-mediated signalling, as suggested by previous studies through real-time parallel NMR analyses (Smith and Ikura, 2014). The differences in the KRas4B^G12V^:Rgl2^RA^ crystal structure (this study) and the Ras:CRAF heterotetramer (Packer *et al*., 2021) provide possible structural explanations for this observation.

## Conclusion

To summarize, our work demonstrates an altered binding kinetics of KRas4B^G12V^ oncogenic mutant with a RalGEF, Rgl2. The G12V mutation resides at the interface of the KRas4B^G12V^-Rgl2^RA^ co-crystal complex. The information may open the way to target oncogenic-*KRAS*-induced tumorigenesis by novel strategies, including interfering molecules for the newly identified interfaces.

## Materials and Methods

### Plasmid constructs

The RA of human Rgl2 was obtained by amplifying a cDNA fragment encoding the position 643 – 740 of the human Rgl2 by PCR with a pair of primers (5’ TACTTCCAATCCATGGGGCCAGGGGCCTCTGATTGCCG3’) and (5’ TATCCACCTTTACTGTCA TGTAGCAGTAGAGGACCTTCGCCGCTGC 3’) using human cDNA prepared from hTERT RPE-1 cells using GoScript Reverse Transcription System (Promega) following the manufacturer’s instruction. The amplified Rgl2^RA^ fragment was cloned into pLEICS2 vector (PROTEX, University of Leicester), which contains a glutathione S-transferase (GST) affinity tag and a Tobacco Etch virus (TEV) potease cleavage site at the N-terminal end of the Rgl2^RA^, using In-Fusion HD EcoDry enzyme (Takara Bio, #638915), following the manufacturer’s instruction.

A cDNA fragment encoding the position 1 – 169 (C-terminal truncated) of the human wildtype KRas4B isoform was amplified by PCR with a pair of primers (5’ TACTTCCAATCCATG ACTGAATATAAACTTGTGGTAGTTGGAGCTG’) and (5’ TATCCACCTTTACTGTCA CTTTTCTTTATGTTTTCGAATTTCTCGAACTAATGTATAG 3’) using human cDNA prepared from hTERT RPE-1 cells as described above. The produced fragment was cloned in pLeics1 plasmid (PROTEX, University of Leicester), which introduces a His6 tag and a TEV cleavage site at the N- terminus. Site directed mutagenesis was conducted to introduce the oncogenic G12V mutation using a pair of DNA oligos (5’ AGTTGGAGCTGTTGGCGTAGGCAAGAGTGCC 3’) and (5’ GTCAAGGCACTCTTGCCTACGCCAACAGCTCCAACTAC 3’) by following the QuikChange^TM^ method (Agilent Technologies, La Jolla, CA, USA).

The RBD of human BRAF was obtained by amplifying a cDNA fragment encoding the position 151 – 232 of the human BRAF by PCR with a pair of primers (5’ TACTTCCAATCCATGTCACCACAAAAACCTATCGTTAGAGTCTTCCTGCC 3’) and (5’ TATCCACCTTTACTGTCAAAGTGGAACATTCTCCAACACTTCCACATGCAATTC 3’) using human cDNA prepared from hTERT RPE-1 cells as described above. The produced fragment was cloned into pLEICS2 vector (PROTEX, University of Leicester) as described above.

The Rgl2^RA^-Halo construct was prepared using the above-described GST-Rgl2^RA^ construct by amplifying the DNA fragment encoding GST-Rgl2^RA^ by PCR using a pair of primers primers (5’ AGGAGATATACATATGTCCCCTATACTAGGTTATTGGAAAATTAAGGG 3’) and (5’ CAGTACCGATTTCGGATCCTGTAGCAGTAGAGGACCTTCGCCG 3’) and inserting the PCR product into pLEICS90 vector (PROTEX, University of Leicester) that adds a Halo tag (Los et al., 2008) at the C-terminal end.

### Protein expression

DE3 Rosetta cells (Novagen) carrying expression plasmids were grown at 37°C in TY media until OD600 reached about 0.6. Then protein expression was induced by adding Isopropyl β-D-1- thiogalactopyranoside (IPTG) to a final concentration of 0.5 mM and keeping the culture at 18°C overnight in a shaking incubator. Cells were collected by centrifugation and resuspended in either Sl1 buffer (Tris 20mM (pH 7.65), NaCl 150 mM, 5 mM imidazole) for His-tagged KRas or in PBS-NaCl buffer (237 mM NaCl, 2.7 mM KCl, 8 mM Na2HPO4, and 2 mM KH2PO4, pH 7.4) for Rgl2^RA^ and BRAF^RBD^. The cell suspensions were then stored at -80°C.

### Protein expression of stable isotope labelling KRas4B^G12V^ and RA of Rgl2 for NMR measurement

The gene encoding human KRas4B^G12V^ and Rgl2^RA^ were constructed into the expression vector pGHL9 and pLEICS2 over-expressed in *Escherichia coli* (*E.coli*) strain BL21 (DE3) and Rosetta, respectively. Uniformly ^13^C, ^15^N-labeled protein was obtained by growing bacteria at 37 °C in M9 minimal media, containing [^13^C6]-glucose and ^15^NH4Cl (Isotec) as the sole carbon and nitrogen source, supplemented with 20 mM MgSO4, 0.1 mM CaCl2, 0.4 mg/ml thiamin, 20 μM FeCl3, salt mix [4 μM ZnSO4, 0.7 μM CuSO4, 1 μM MnSO4, 4.7 μM H3BO3], and 50 mg/L ampicilin. KRas4B^G12V^ NMR sample was prepared essentially as described previously (Ito *et al*., 1997). Protein expression of Rgl2^RA^ was induced by adding 119mg/L IPTG at an OD 600 nm of 0.5. After 18 h of further growth, cells were harvested, and washed with a pH 7.5 lysis buffer [50 mM Tris-HCl, 25% sucrose, and 0.01% NP-40]. Uniformly ^15^N-labeled KRas4B^G12V^ and Rgl2^RA^ were produced by the identical steps unless growing cells in M9 medium containing [^12^C6]-glucose and ^15^NH4Cl (Isotec).

### Purification of GST-tagged Rgl2^RA^, Rgl2^RA^-Halo and BRAF^RBD^

Bacteria cell suspensions were thawed and supplemented with Triton X-100 to a final concentration of 0.1 % (v/v). Cells were broken by a probe sonicator and insoluble materials were removed by centrifugation. The supernatant was mixed with glutathione (GSH) beads (GE Healthcare, #17-0756-01) and incubated for 20 min at 4°C. The GSH beads were washed three times with the PBS-NaCl buffer. The GST-Rgl2^RA^ and GST-BRAF^RBD^ fusion proteins were eluted by the elution buffer (50mM Tris-Cl (pH 8.0), 100mM NaCl, 5mM GSH) for BLI experiments. The Rgl2^RA^ and Rgl2^RA^-Halo were separated from the GST tag through cleavage by TEV protease, which was prepared as previously described (Kapust et al., 2001; Tropea et al., 2009), for structural analysis and mass photometry analysis. The obtained Rgl2^RA^, Rgl2^RA^-Halo or GST- Rgl2^RA^/GST-BRAF^RBD^ fusion proteins were concentrated using a concentrator (10 KDa MWCO, Merck, Amicon Ultra centrifugal filters, #UFC901024) and filtrated using a centrifugal filter unit (Milipore, Ultrafree, #UFC30GV00) before conducting size exclusion chromatography (SEC) with a gel-filtration column (GE Healthcare, HiLoad Superdex 75) attached to an FPLC system. SEC was carried out in the gel filtration (GF) buffer (20 mM Tris-Cl (pH 7.65), 100 mM NaCl, 5mM MgCl2, 1mM Tris (2-carboxyethyl) phosphine (TCEP)).

### Purification of His-tagged KRas4B wildtype and oncogenic G12V mutant proteins

Bacteria cell suspensions were supplemented with Triton X-100 to a final concentration of 0.1 % (v/v). Cells were broken by a probe sonicator and insoluble materials were removed by centrifugation. The soluble cell lysates were applied on a Ni-sepharose excel (GE Healthcare, #17-3712-01), packed in a column of 4 ml bed volume with Sl1 buffer buffer (20mM Tris, pH6.5, 150mM NaCl, 5 mM imidazole). The column was washed with 20 ml of SL1 Buffer, then with 20 ml of Sl3 Buffer (20mM Tris, pH6.5, 150mM NaCl, 6mM Imidazole) and finally with 15 ml of Sl4 Buffer (20mM Tris, pH6.5, 150mM NaCl, 10mM Imidazole). The His-tagged KRas protein was eluted from the column by applying 10 ml of Elution buffer (50mM Tris, pH 7.65, 150mM NaCl, 200mM Imidazole), followed by 10 ml of 1 M imidazole. In order to remove the His6-tag at the N-terminal end, TEV protease, prepared as previously described (Tropea *et al*., 2009), was added to the elution fraction to about a 2 % molar ratio of the His6-KRas4B preparation and incubated overnight at 4°C. The cleaved KRas samples were further purified by SEC using a gel-filtration column (GE Healthcare, HiLoad Superdex 75) in the GF buffer (20 mM Tris-Cl (pH 7.65), 100 mM NaCl, 5mM MgCl2, 1mM Tris (2-carboxyethyl) phosphine (TCEP)). The purified KRas (WT) and KRas(GV) concentrations were determined by absorbance at 280 nm. The extinction coefficient (ε) for KRas (WT) and KRas(GV) was estimated to be 19685 cm-^1^ *M* ^-1^, by taking into account that the bound GTP adds 7765 cm-^1^ *M* ^-1^ (Smith and Rittinger, 2002), and the molecular weights (including GTP) were estimated to be 19856 and 19898, respectively. Nucleotide exchange of the purified KRas4B wildtype or G12V proteins was carried out essentially as previously described (Ito *et al*., 1997). The proteins were diluted 10 times by the exchange buffer (20mM Tris-Cl (pH7.5) 1mM EDTA, 1mM TECP), and the sample was supplemented with EDTA to a final concentration of 5 mM. The sample was mixed with about 10 time molar excess of guanosine 5′- [β,γ-imido]triphosphate (GMMPPNP, SIGMA G0635) or guanosine 5’-triphosphate (GTP, SIGMA G9977) or guanosine 5’ –diphosphate (GDP, SIGMA G7127). The reaction was incubated at 37℃ for 20 minutes then put it on ice for 20 minutes. The protein was terminated by adding ice cold MgCl2 to a final concentration of 20 mM. The excess nucleotides were removed by SEC as described below.

### Purification of isotope labelling KRas4B^G12V^

All the procedures described below were carried out at 4 °C unless otherwise stated. All the isotope-labelled KRas4B^G12V^ samples were purified by the same step. The cells dispersed in the lysis buffer was disrupted by sonication for 30 min on ice with hen egg lysozyme (0.1 mg/mL). The cell debris was clarified by centrifugation at 14,000 g for 1 h. The supernatant was loaded onto a 25 mL of DEAE-Sepharose Fast Flow (Cytiva) anion exchange equilibrated with buffer A [50 mM Tris-HCl (pH 7.5), 1 mM MgCl2, 1 mM dithiothreitol (DTT), 0.1 mM APMSF (FUJIFILM Wako)]. After washing the column with buffer A until sufficiently low of UV absorption at 280 nm, the KRas4B^G12V^ protein was eluted by linearly increasing the concentration of KCl from 0 to 350 mM with a flow rate of 0.5 mL/min in buffer A. The fractions containing the target protein were concentrated to 5 mL with Amicon Ultra-15 10 kDa (Merck). The concentrated sample was loaded onto a 320 mL of HiLoad Superdex 75 (GE Healthcare Life Science) gel filtration with a flow rate of 0.8 mL/min using FPLC systems (AKTA pure 25, GE Healthcare Life Science). The 5 mL sample concentrated from the fractions involving the target proteins with Amicon Ultra-15 10 kDa was loaded on Resource Q (GE Healthcare Life Science) anion-exchange column equilibrated with buffer A using the FPLC systems. After washing the column with 30 mL of buffer A, KRas4B^G12V^ was eluted by an KCl, the KRas4B^G12V^ protein was eluted by linearly increasing the concentration of KCl from 0 mM to 350 mM with a flow rate of 1 mL/min in buffer A. The purification of isotope labelled Rgl2^RA^ was performed by the same step described above. The purity of the KRas4B^G12V^ and Rgl2^RA^ samples in each step was confirmed by SDS-PAGE. Protein concentrations were determined by NanoDrop 2000 (ThermoFisher) measuring UV absorption at 280 nm. KRas4B^G12V^ samples for NMR measurements were concentrated and dissolved in NMR buffer A (90% ^1^H2O/ 10% ^2^H2O containing 20 mM Tris-HCl (pH 7.5), 100 mM NaCl, 5 mM MgCl2, 1 mM β-mercaptoethanol). Rgl2^RA^ samples for NMR measruments were NMR buffer B (90% ^1^H2O/ 10% ^2^H2O containing 1 mM Na2HPO4-NaH2PO4 (pH 7.4), 150 mM NaCl).

### KRas4B-Rgl2^RA^ and KRas4B-BRAF^RBD^ binding measurements using BLI

Octet R8 (Sartorius) was used for Biolayer interferometry assays of KRas4B (G12V or WT) and Rgl2^RA^ and BRAF^RBD^ interactions. Anti-GST biosensors (Sartorius #18-5096) were used to immobilise GST-Rgl2^RA^ (provided as a 0.7 µM solution in the binding reservoir well, the concentration was determined by absorbance at 280 nm using the following values; the extinction coefficient (ε) 48820 cm-^1^ *M* ^-1^, and the molecular weight 37495.18), GST-BRAF^RBD^ (encoding BRAF a.a. 151-232, N-terminally fused with GST, provided as a 0.7 µM solution in the binding reservoir well, the concentration was determined by absorbance at 280 nm using the following values; the extinction coefficient (ε) 56840 cm-^1^ *M* ^-1^, and the molecular weight 36024.95). The baseline was stabilised in GF buffer (20mM Tris pH7.65, 100 mM NaCl, 5mM MgCl2, 1mM TCEP) for 200sec. As a negative control, GST only (provided as a 0.7 µM solution in the binding reservoir well) was used. The association of KRas4B was measured for 400sec in 5 or 6 serial dilutions concentrations. For Rgl2^RA^ binding, the KRas concentrations ranged from 200 nM - 15 µM, and for BRAF^RBD^ binding, the KRas concentrations ranged from 20 nM – 5 µM. The dissociation steps were measured in fresh GF buffer for 400sec. For each assay a biosensor immobilised with GST only and a sample well with only buffer instead of KRas4B (WT/G12V) was set up for double referencing. The experiments were conducted at 20°C. The resulting data were processed using Octet Analysis Studio (ver. 13.0)(Sartorius) and reference biosensor (loaded with GST-only) and reference wells (containing no KRas) were subtracted from sample wells (double reference).

### Analysis of bound nucleotide

The nucleotide-binding status of KRas4B^WT^ and KRas4B^G12V^ were examined by denaturating the proteins and detecting the released nucleotides, essentially following the previous studies (Smith and Rittinger, 2002). About 2 nmoles of KRas4B molecules were adjusted to the volume of 200 µl with GF buffer. Add 12.5 µl of 10% perchloric acid to denature and precipitate the protein. The supernatant was neutralised either by 8.75 µl of 4M CH3COONa, pH 4.0, or 12 µl of 1M Tris-Cl (pH 8.8). The sample was centrifuged again, and the supernatant was analysed by High-performance liquid chromatography (HPLC) using an ion exchange column (Partisil 10 SAX column, Whatman). For the result presented in Fig. 1A, the chromatography was run using 0.6 M NH4H2PO4 (buffer B) at a flow rate of 0.8 ml/min. For the result presented in Fig. 1D, the column was first equilibrated with 10 mM NH4H2PO4 (buffer A), and the column was run with the following gradient condition with 0.6 M NH4H2PO4 (buffer B) at a flow rate of 0.8 ml/min. Step1: 100% buffer A (0% buffer B) for 11 min, Step2: a gradient increase to 40% buffer B over 6 min, Step3: a gradient increase to 50% buffer B over 23 min, Step4: a gradient increase to 100 % buffer B over 1 min, Step 5: 100 % buffer B for 19 min, Step 6: a gradient decrease to 0% buffer B (100 % buffer A) over 1 min, and Step 7: 100% Buffer A for 14 min. The nucleotides were detected by 254 nm absorption. As a reference control, 1 µl of 10 mM GTP or GDP was diluted to 200 µl GF buffer and was processed in the same manner as protein samples.

### Crystallography

The purified and GMPPNP-loaded KRas4B (G12V) and Rgl2^RA^ were mixed in the GF buffer and the complex was purified on SEC using a gel-filtration column (GE Healthcare, HiLoad Superdex 75). The peak fractions containing both proteins in the 1:1 ratio were collected and concentrated to set up crystallization screenings. Crystals of KRas4B (G12V) and Rgl2^RA^ were obtained using sitting-drop vapour diffusion at room temperature, with 100nl of protein (11.6mg/ml) against 100nl of crystallisation buffer (0.2M sodium/potassium phosphate ph 7.5, 0.1M HEPES pH 7.5, 22.5% v/v PEG smear Medium (Molecular Dimensions MD2-100-259), 10% v/v glycerol. The crystals were frozen in liquid nitrogen with 20% glycerol as cryoprotectant. Data were collected at Diamond beamline I04. AIMLESS (Evans and Murshudov, 2013) was used for data reduction before obtaining phaser solution using the HRas-RALGDS^RA^ complex structure (PDB ID 1LFD) as search model with PhaserMR (McCoy et al., 2007). The structure was built using multiple rounds of refinements using PDBredo, REFMAC, PHENIX and COOT (Emsley et al., 2010; Joosten et al., 2014; Murshudov et al., 2011; Torices and Munoz-Pajares, 2015). The coordinates of the complex have been deposited to the Protein Data Bank (PDB) under access code 8B69.

### Circular dichroism (CD) spectroscopy

KRas4B(WT) and (G12V) proteins at a concentration of 20 μM were prepared in the CD buffer (50 mM phosphate (pH 7.6), 1.5 mM Tris (pH 7.6), 5mM MgSO4, 7.5 mM NaCl, 0.375 mM MgCl2), placed in a quartz cuvette of 0.1 cm path length and CD spectra were recorded at wavelengths ranging from 195 to 250 nm using a Chirascan^TM^-plus CD spectrometer (Applied Photophysics) at 20 °C. The melting curves of these proteins were examined at 220 nm at temperatures ranging from 20°C to 90°C. Measurements were conducted at 1 °C increment.

### NMR spectroscopy

KRas4B^G12V^ NMR sample was prepared essentially as described previously (Ito *et al*., 1997). The bacteria expression plasmid for HRas was modified to encode KRas4B^G12V^ by site-directed mutagenesis and used to produce KRas4B^G12V^. Loading of a GTP analogue, GMPPNP (Jena Bioscience) was conducted essentially as previously described (Ito *et al*., 1997). Rgl2^RA^ sample was prepared as described above. All NMR samples were measured in 20 mM Tris-Cl (pH 7.65), 100 mM NaCl, 5mM MgCl2, and 0.1% β-mercaptoethanol at 303 K. All spectra were analysed with the CcpNmr Analysis 2.5.1 software. Backbone chemical shifts of KRas4B^G12V^ and Rgl2 have been deposited to the BioMagResBank with accession ID 34754 and Protein Data Bank with accession ID 8AU4, respectively.

All NMR experiments were performed at 25°C probe temperature in a triple-resonance cryoprobe fitted with a z-axis pulsed field gradient coil, using Bruker AVANCE-III HD 600 MHz spectrometers. All spectra were processed with the Azara software package (Boucher). For the 3D data, the two-dimensional maximum entropy method (2D MEM) or Quantitative Maximum Entropy (QME) (Hamatsu et al., 2013) were applied to obtain resolution enhancement for the indirect dimensions. All NMR spectra were visualized and analyzed using the CcpNmr Analysis 2.5.0 software (Vranken et al., 2005). All of the 3D triple-resonance experiments used for the assignments of KRas4B and Rgl2^RA^ were performed on ^13^C/^15^N samples in NMR buffer A and B, respectively. The backbone ^1^H^N^, ^13^C’, and ^15^N for KRas4B and Rgl2^RA^, and side-chain ^13^C^μ^ and ^13^C^μ^ resonance assignments for Rgl2^RA^ were achieved by analyzing six types of 3D triple-resonance experiments, HNCO, HN(CA)CO, HNCA, HN(CO)CA, CBCANNH, and CBCA(CO)NNH. 3D HBHA(CBCACO)NH, H(CCCO)NH, (H)CC(CO)NH, HCCH-COSY, and HCCH-TOCSY experiments on the ^13^C/^15^N-labeled Rgl2^RA^ were performed for side-chain ^1^H and ^13^C resonance assignments. A 15 ms ^13^C isotropic mixing time was employed for the (H)CC(CO)NH, H(CCCO)NH and HCCH-TOCSY experiments. For the collection of NOE-derived distance restraints of Rgl2^RA^, 3D ^15^N- separated and 3D ^13^C-separated NOESY-HSQC spectra were measured on the ^13^C/^15^N-labeled Rgl2^RA^. A 100 ms NOE mixing period was employed for the 3D NOESY experiments. All 2D and 3D NMR data were recorded using the States-TPPI protocol for quadrature detection in indirectly observed dimensions. Water flip-back ^1^H pulses and the WATERGATE pulse sequence were used for solvent suppression in the experiments performed on ^15^N-labeled, and ^13^C/^15^N-labeled samples, whereas presaturation and gradient-spoil pulses were used for ^13^C-labeled samples.

A series of 2D ^1^H-^15^N HSQC spectra were measured for titration experiments of ^15^N-labelled KRas4B^G12V^ with in the presence of non-labelled Rgl2^RA^. The experiments were performed in the NMR buffer at 25 °C and the peptide concentration was increased stepwise (for the ^15^N- KRas4B^G12V^ / Rgl2^RA^, its molar ratio of 1:0.25, 1:0.5, 1:0.75, 1:1, 1:1.5, and 1:2 were used, while for the ^15^N-Rgl2^RA^ / KRas4B^G12V^, its molar ratio of 1:0.25, 1:0.5, 1:1, 1:1.5, 1:2, and 1:3 were used). The mean chemical shift difference Δδave for each amino acid was calculated as [(Δδ^1^H^N^)^2^ + (Δδ^15^N)^2^]^1/2^ where Δδ^1^H^N^ and Δδ^15^N are the chemical shift differences (Hz) between KRas4B^G12V^ or Rgl2^RA^ on their own and the proteins in the presence of the other side.

### NMR structure calculation

Intra-residual and long-range NOEs were automatically assigned by the program CYANA with the use of automated NOE assignment and torsion angle dynamics (Güntert and Buchner, 2015). The peak position tolerance was set to 0.03 ppm for the 1H dimension and to 0.3 ppm for the 13C and 15N dimensions. Hydrogen-bond and dihedral angle restraints were not used. CYANA calculations were started from 100 conformers with random torsion angle values, simulated annealing with 50,000 torsion angle dynamics steps was applied. The 20 conformers with the lowest final target-function values of CYANA were selected and optimised with OPALp 2.1 [22,23] using the AMBER force field [24,25]

### NMR relaxation experiment

The longitudinal relaxation times (*T*1), the transverse relaxation times (*T*2), and the steady-state heteronuclear {^1^H}-^15^N NOEs were measured at 25 °C using uniformly ^2^H/^13^C/^15^N-labeled KRas4B^G12V^ with non-labelled Rgl2^RA^ on a Bruker Avance III HD 600 MHz spectrometer equipped with a cryogenic H/C/N triple-resonance probe head. Each experiment was acquired in a pseudo 3D manner. 8 relaxation delays in the range 20–1800 ms and 8 delays between 8.7–139 ms were used for the *T*1 and *T*2 experiments, respectively. NOE ratios were obtained from intensities in experiments recorded with (1 s relaxation delay followed by 10 s saturation) and without (relaxation delay of 11 s) saturation. The overall rotational correlation time, effective correlation times, and order parameters were obtained by the program relax 5.0. with spectral density functions as defined by the Lipari-Szabo model-free approach. A theoretical rotational correlation time (τ*c*) was calculated by the Stokes-Einstein-Debye relation,

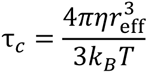

Where *η* is the viscosity, *r*eff is the effective hydrodynamic radius of a molecule, *k*B is the Boltzmann constant, and *T* is the temperature. The rotational correlation time of the KRas4B^G12V^:Rgl2^RA^ complex was estimated with *T* = 298.0 K, *η* = 0.890 mPa, and *r*eff = 27.1 Å.

### Mass photometry measurement

For KRas4B^G12V^ and Rgl2^RA^ complex formation, freshly purified KRas4B^G12V^ and Rgl2^RA^ were mixed in GF buffer, concentrated to about 500 µl using a concentrator (Merck, Amicon Ultra centrifugal filters, #UFC901024), and loaded to a gel-filtration column (GE Healthcare, HiLoad Superdex 75). The peak fractions containing both proteins in the 1:1 ratio were collected and the concentration was estimated by OD280 and the predicted extinction coefficient of 24155 M^-1^cm^-1^. The sample was diluted to 40 nM in the GF buffer and a 20 µL aliquot was subjected to mass photometry immediately after the dilution (OneMP, Refeyn). For KRas4B^G12V^ and Rgl2^RA^- Halo complex formation, both proteins were separately purified by SEC as described above and mixed at the 1:1 ratio to generate a premix 2 µM sample, which was further diluted to 20 nM, 50 nM, 100 nM and 250 nM before the measurement.

The measurement was done in conventional microscope cover glass (Marienfeld, n° 1.5H) cleaned by rinsing with deionized water (′5) and isopropanol (′5) followed by drying under a N2 flow, using a silicon gasket (Grace Bio-labs) in order to confine the sample. Adsorption of individual molecules of complex was detected across an imaging area of 10.8 µm by 2.9 mm.

Video recordings from interferometric scattering microscopy for a duration of 60s were obtained and the single events corresponding to surface adsorption of the complex were identified using AcquireMP software (Refeyn). Data analysis was performed using DiscoverMP software (Refeyn) and OriginPro 2021 (OriginLab).

### Graphical representation of protein structures

Protein structure images were generated using Pymol (The PyMOL Molecular Graphics System, Version 1.2r3pre, Schrödinger, LLC). Electrostatic surface charge potential images were produced using Pymol vacuum electrostatics function. Amino acid residues in the interaction surfaces of protein complexes in a PDB format were predicted using LIGPLOT (Wallace et al., 1995).

## Acknowledgments

We thank the help and expert advice from the University of Leicester colleagues Peter Moody, Thomas Schalch, Ian Eperon, Mohammed Bhogadia, Luke Bailey, Kyle Daynes, Idir Malki, and Ziad Ibrahim. Biolayer interferometry experiments were conducted with the guidance of Holly Birchenough and Thomas Jowitt (Biomolecular Analysis Core Facility, University of Manchester).

We gratefully acknowledge financial support from Wellcome Trust ISSF COVID career support and inclusion scheme 204801/Z/16/Z, University of Leicester MSc program (to A.H., M.A.M.C. and K.T.), University of Leicester College PhD programme (to M.T., L.R.A. and K.T.), BBSRC MIBTP PhD programme BB/T00746X/1 (to S.M. and K.T.), the Funding Programs for Core Research for Evolutional Science and Technology (CREST; JPMJCR13M3 to Y.I., JPMJCR21E5 to T.I.) the Japan Science and Technology Agency (JST), Grants-in-Aid for Scientific Research (JP15K06979 to T.I., JP19H05645 to Y.I.) and Scientific Research on Innovative Areas (JJP15H01645, JP16H00847, JP17H05887 and JP19H05773 to Y.I. P26102538, JP25120003, JP16H00779 and JP21K06114 to T.I.) from the Japan Society for the Promotion of Science (JSPS), Shimazu foundation, and the Precise Measurement Technology Promotion Foundation.

## Author Contributions

M.T., T.I., Y.I., C.D. and K.T. designed research. M.T., T.I., N.T., S.K., S.M., L.R.A, A.H., C.B.-A., S.S., M.A.M.C., C.D. and K.T. performed research. M.T., T.I., N.T., L.F., S.K., C.B.-A., S.S., B.R.-A, A.J.H., Y.I., J.W.R.S., C.D. and K.T. analyzed data. M.T., T.I., C.D. and K.T. wrote the manuscript with feedback from all the authors.

## Classification

Biochemistry, Structural Biology

## Legends to Figuress

**Figure S1.**
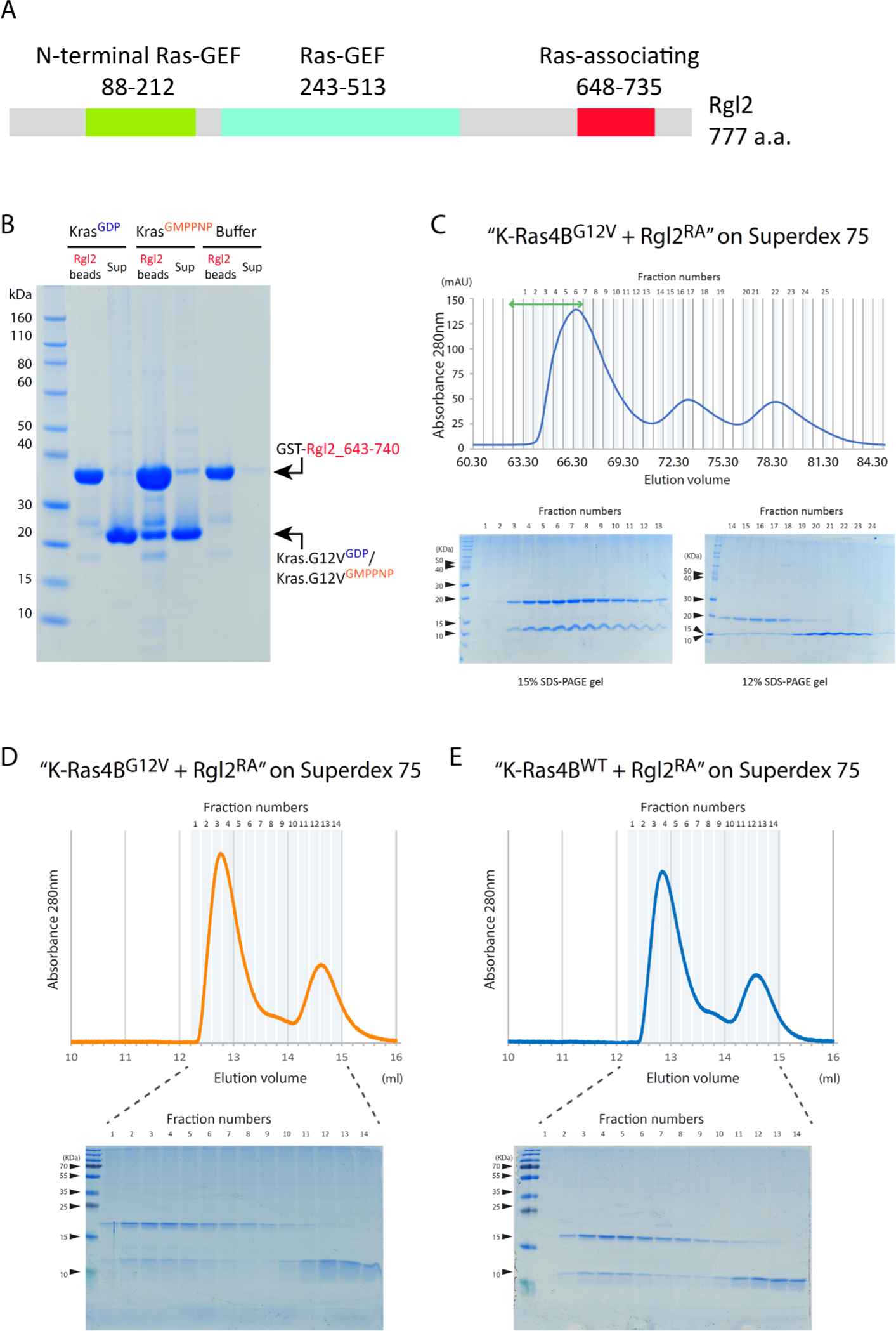
Rgl2 Ras binding domain interacts with active KRas4B^G12V^. (A) A schematic diagram of the domain organisation of human Rgl2 as defined in the UniProt database (Uniprot number: O15211). A domain spanning amino acid residues 648-735 is annotated as “Ras-associating” by PROSITE annotation rule PRU00166. In this work, we call this domain Rgl2^RA^. (B) Rgl2^RA^ interacts with active KRas4B^G12V^. Bacteria recombinant Rgl2^RA^ fragment spanning the amino acid residues 643-740 of Rgl2 was fused with GST, and the fusion protein was fixed on glutathione beads. Recombinant KRas4B^G12V^ 1-169, loaded with either GDP or non-hydrolysable GTP analogue, GMPPNP, was applied on the beads to examine the protein-protein interaction. GST-Rgl2^RA^ interacted only with the GMPPNP-loaded KRas4B^G12V^. (C) KRas4B^G12V^:Rgl2^RA^ complex was purified using size exclusion chromatography. The fractions were analysed by SDS-PAGE gels (lower panel) according to the elution profile (upper panel). Fractions indicated by the green double-arrow line were used for crystallisation trials. (D) and (E) The size exclusion chromatography elution profiles of the KRas4B-Rgl2^RA^ complex. Rgl2^RA^ and KRas4B^G12V^ (D) or KRas4B^WT^ (E) were mixed at the 3:1 molar ratio and applied to Superdex 75 10/300 GL. The elution profiles (upper panels) show little difference between the two samples. Peak fractions were analysed by SDS-PAGE gel (lower panel).

**Figure S2.**
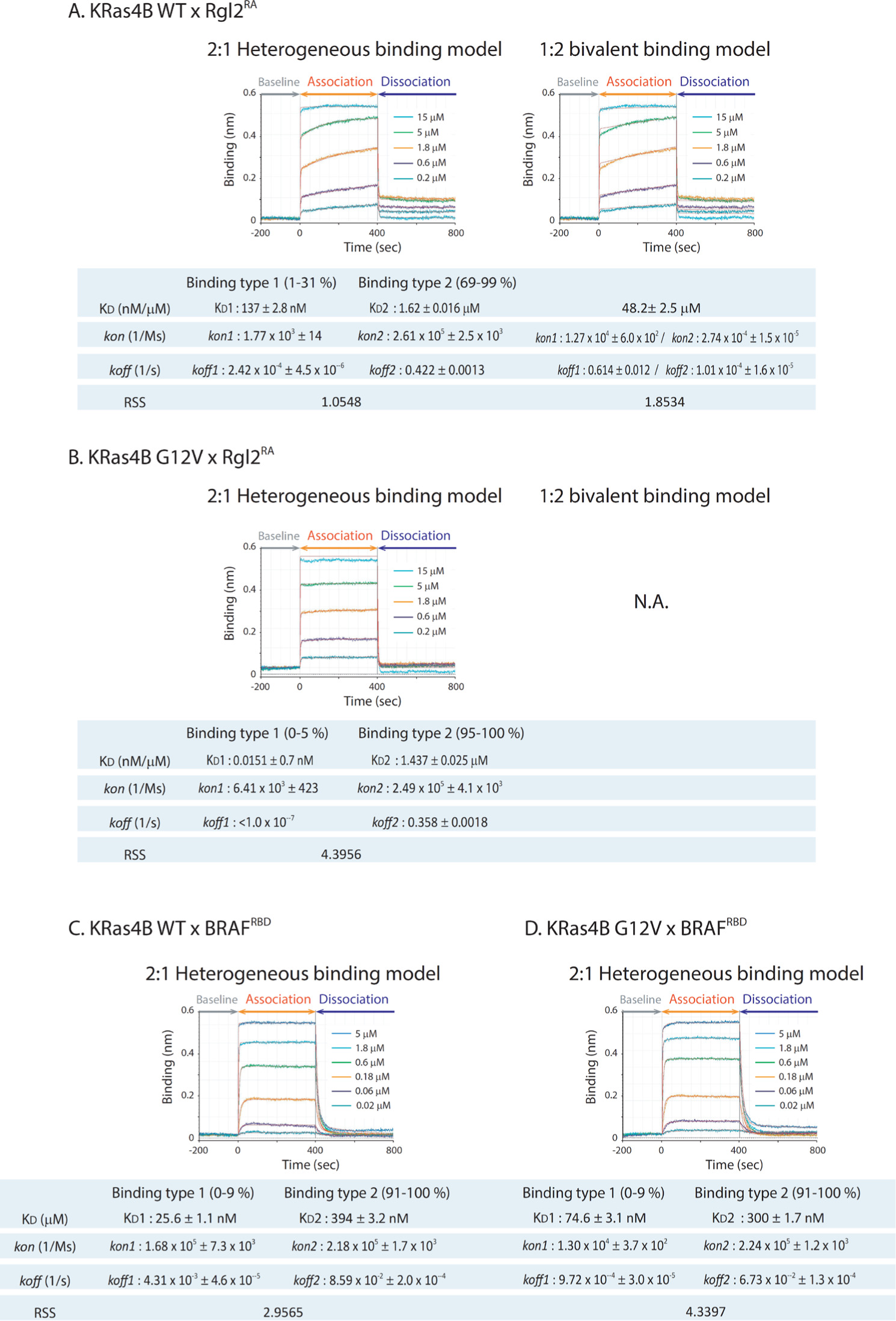
Analysis of KRas4B-effector binding kinetics. (A) and (B) Analyses of binding kinetics of GST-Rgl2^RA^ and KRas4B^WT^ or KRas4B^G12V^. The BLI sensorgram data presented in Fig. 1B for the binding kinetics between Rgl2^RA^ and KRas4B^WT^ (A) or KRas4B^G12V^ (B) were fitted for the 2:1 heterogeneous binding and the 1:2 bivalent binding models using Octet Analysis Studio 13.0 (Sartorius). The fitted curves are shown in red, and the deduced KD, *kon*, *koff*, and the residual sum of squares (RSS) are shown below the sensorgrams. For the 2:1 heterogeneous binding model, the values for the two binding types (Binding type 1 and 2) are shown. The proportions of each Binding type (1 or 2) varied among the samples of different concentrations; hence, the varied percentage ranges are indicated. For the 1:2 bivalent binding model, the KRas4B^G12V^ data could not be fitted. (C) Analyses of binding kinetics of GST- BRAF^RBD^ and KRas4B^WT^ or KRas4B^G12V^. The BLI sensorgram plots presented in Fig. 1C were fitted for the 2:1 heterogeneous binding model using Octet Analysis Studio 13.0 (Sartorius). The fitted curves are shown in red, and the deduced KD, *kon*, *koff* and RSS are shown below the sensorgrams. The proportion of each binding type (Binding type 1 or 2) varied among the samples of different concentrations; hence, the varied percentage ranges are indicated.

**Figure S3.**
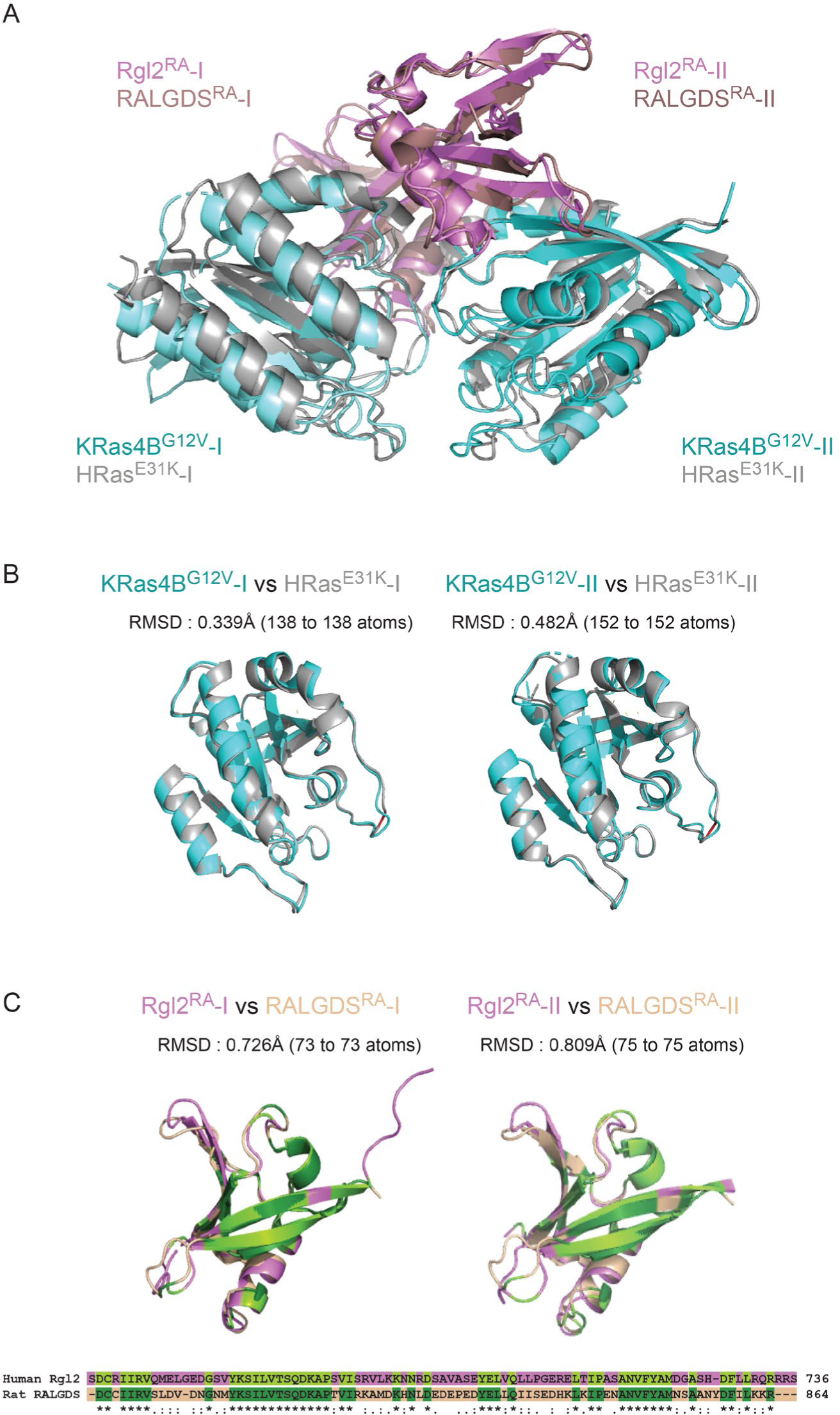
KRas4B^G12V^:Rgl2^RA^ complex is highly similar to HRas^E31K^:RALGDS^RA^ complex. (A) The structures of the KRas4B^G12V^:Rgl2^RA^ complex and the HRas^E31K^:RALGDS^RA^ complex (PDB: 1LFD) were superimposed by aligning the RA chains. (B) and (C) Superimposition of each corresponding chain of the KRas4B^G12V^: Rgl2^RA^ complex and HRas^E31K^:RALGDS^RA^ complex individually shows that each chain is similar to its respective counterpart structure. (B) KRas4B^G12V^ and HRas^E31K^ chains are in cyan and grey. (C) Rgl2^RA^ and RALGDS^RA^ are in purple and light pink. Amino acid sequence similarities of human Rgl2^RA^ and rat RALGDS^RA^ are shown below the RA cartoon representations, and the conserved amino acids in Rgl2^RA^ and RALGDS^RA^ are shaded in light green and dark green, respectively. The structural similarity between Rgl2^RA^ and RALGDS^RA^ was seen beyond the regions composed of the conserved amino acids. The RALGDS^RA^ sequence is derived from PDB 1LFD, which is rat RALGDS^RA^, and the numbering was done according to Uniprot Q03386. Root-mean-square deviation of atomic positions (RMSD) values were generated using Pymol.

**Figure S4.**
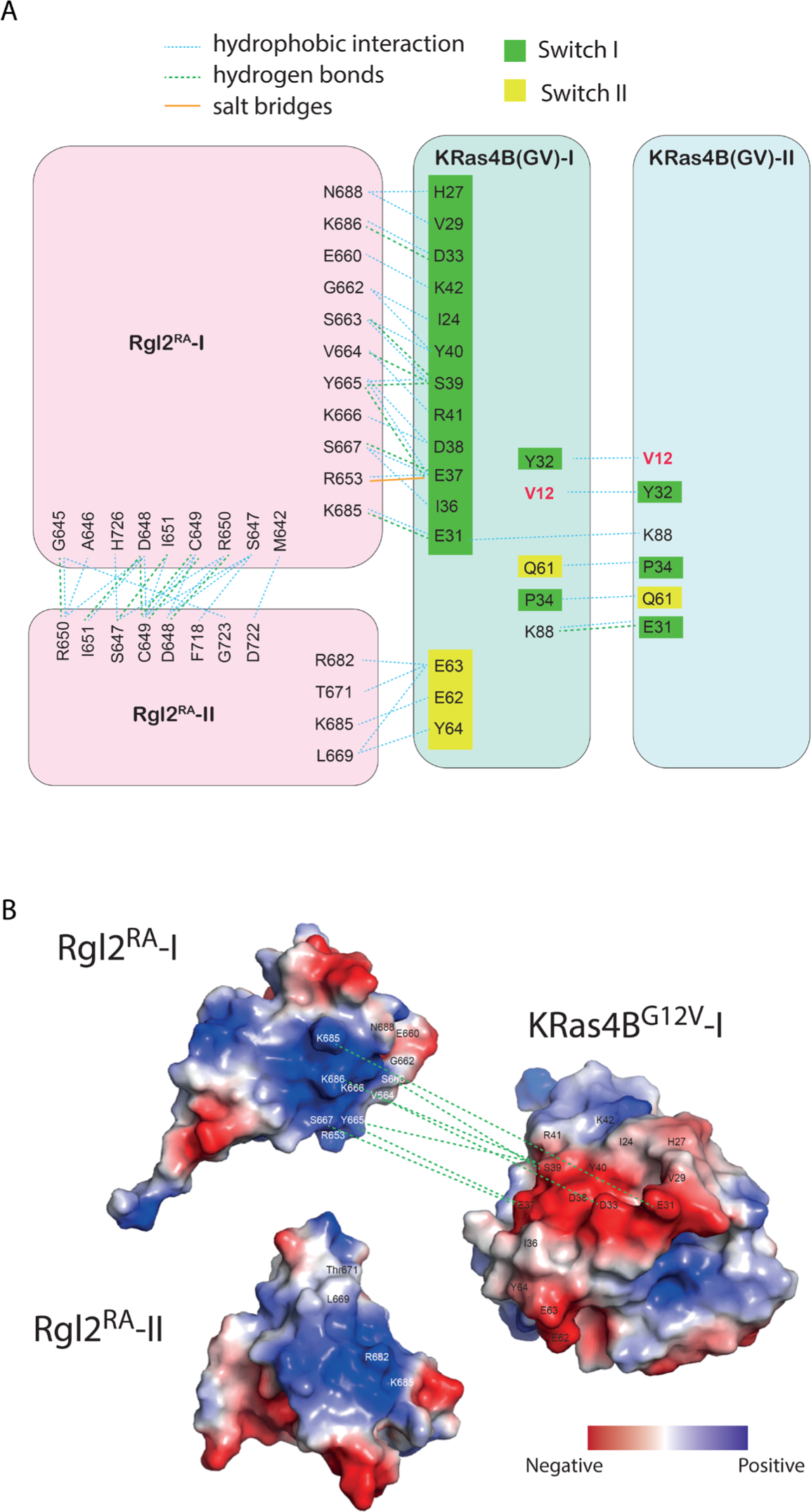
Schematic representation of the intermolecular contacts observed in the X-ray structure. (A) The summary of the diagrams presented in Fig. 3. The residues in KRas4B^G12V^ Switch-I region are shown in green, in the Switch-II region are shown in the yellow background, and those for Rgl2^RA^ molecules 1 and 2 are shown in pink. V12 oncogenic mutation is in red. Hydrogen bonds, salt bridges and hydrophobic interactions were predicted by LIGPLOT and represented by green dashed lines, solid orange lines and blue dotted lines, respectively. (B) The electrostatic surface charge shows that the negatively charged Switch-I and Switch-II regions of KRas4B^GV^ interact with the positively charged surface of Rgl2^RA^-1 and Rgl2^RA^-2. Surface charge potential was computed using the Pymol vacuum electrostatics function, and the negatively charged and positively charged areas are shown in red and blue, respectively.

**Figure S5.**
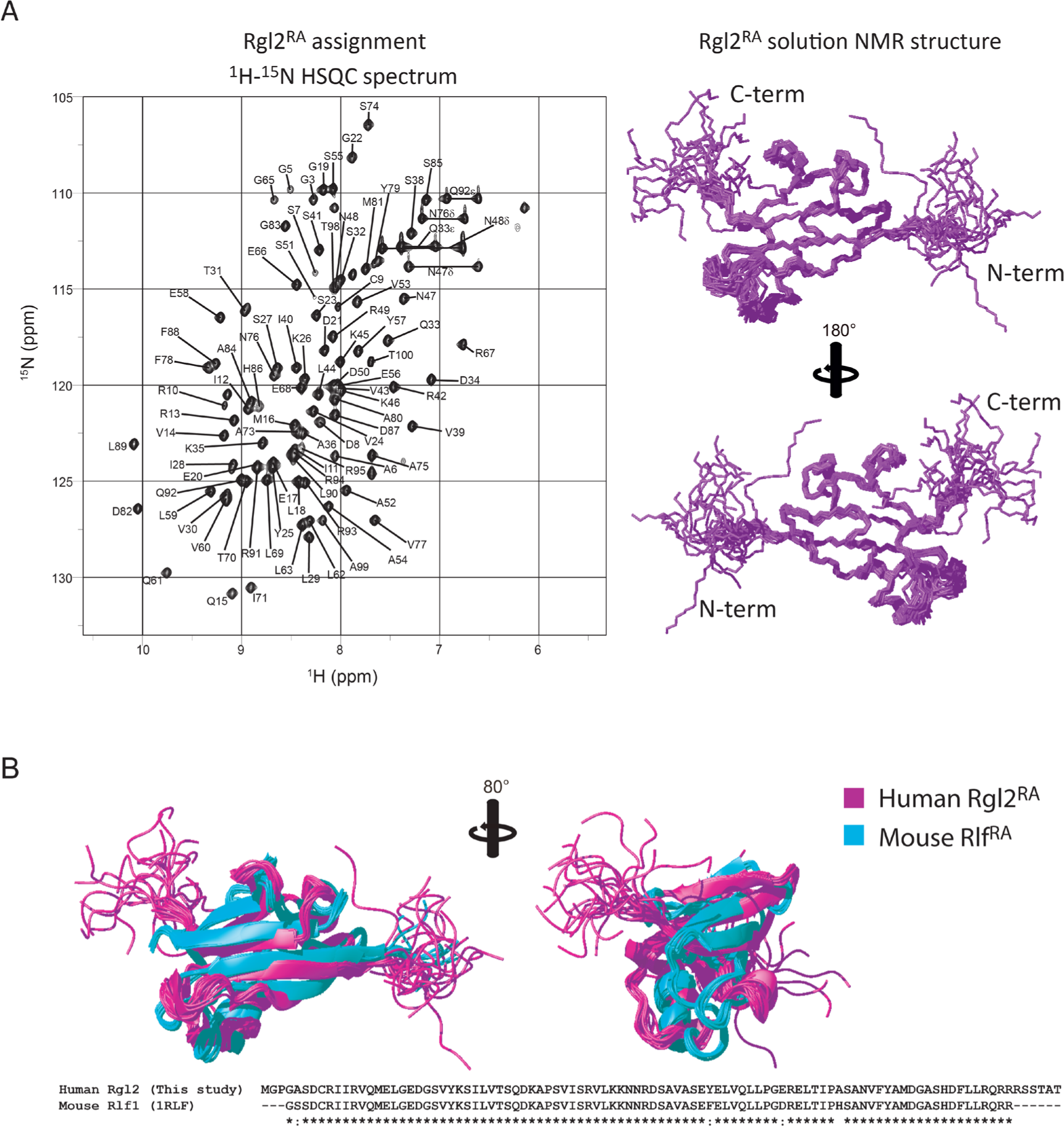
KRas4B^G12V^:Rgl2^RA^ complex formation in solution. (A) Rgl2^RA^ solution structure determined by NMR. The 2D ^1^H-^15^N-HSQC spectrum of the assigned residues of the Rgl2^RA^ is shown (left). The Rgl2^RA^ retains the ββαββαβ ubiquitin-fold structure (right). 20 Rgl2 NMR structures are superimposed. (B) Comparison between the human Rgl2^RA^ and mouse Rlf^RA^ (PDB 1RLF) solution structures. Rgl2^RA^ is presented in magenta, and 20 NMR structures are superimposed. Rlf^RA^ is in cyan, and 10 NMR structures are superimposed. The primary structures of the two constructs are listed below.

**Figure S6.**
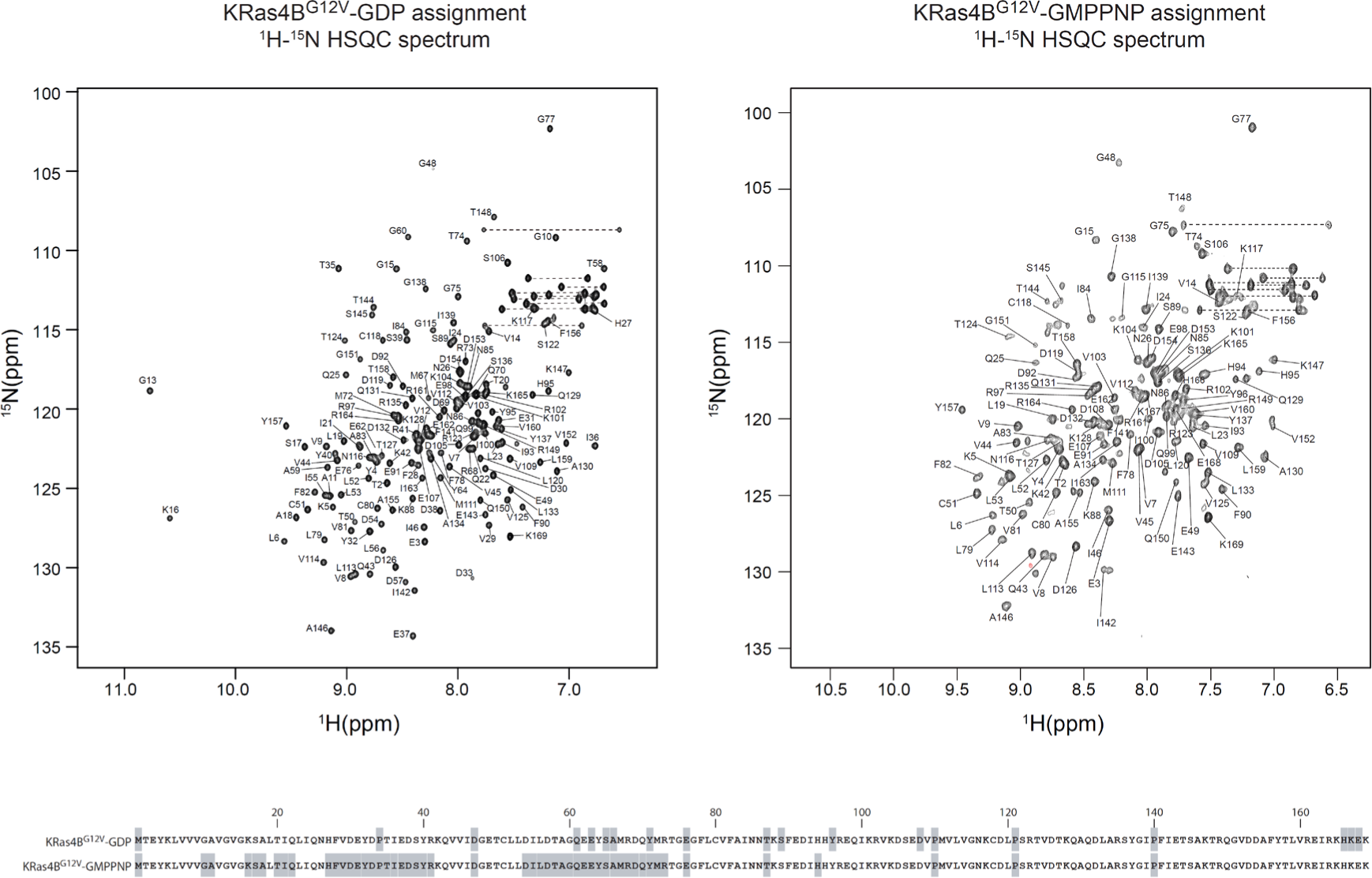
Backbone resonance assignment of GDP-bound and GMPPNP-bound KRas4B^G12V^. The 2D ^1^H-^15^N-HSQC spectra of GDP-Bound and GMPPNP-bound KRas4B^G12V^. Cross peaks are labelled with their corresponding backbone assignments. Residues which could not be assigned are shaded in grey in the primary sequence shown at the bottom.

**Figure S7.**
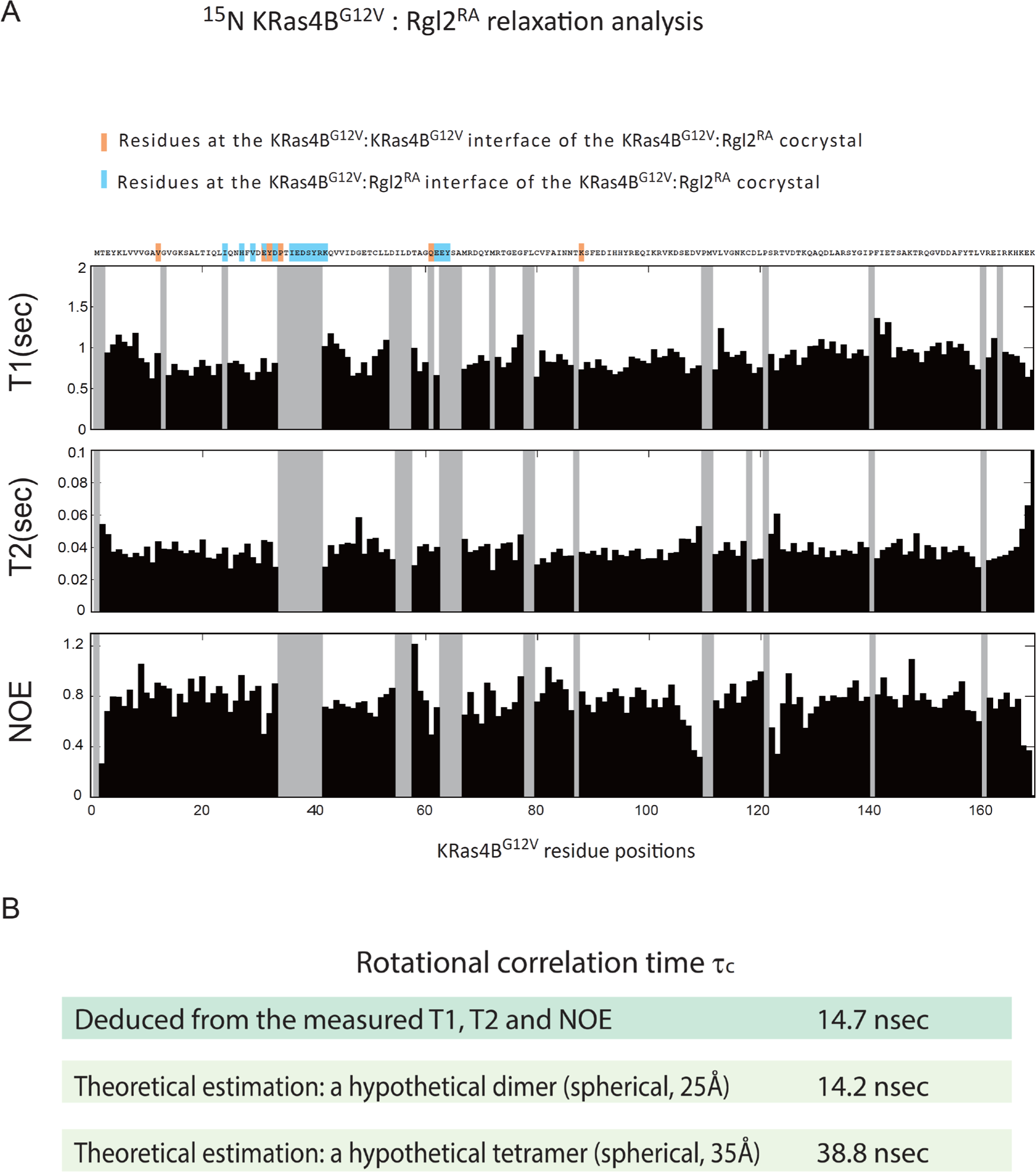
*T*1, *T*2, and {^1^H}-^15^N NOE for backbone ^15^N resonances of ^15^N-KRas4B^G12V^:Rgl2^RA^ complex. (A) The longitudinal relaxation times (*T*1), the transverse relaxation times (*T*2), and the steady-state heteronuclear {^1^H}-^15^N NOEs were measured using uniformly ^2^H/^13^C/^15^N-labeled KRas4B^G12V^ with non-labelled Rgl2^RA^. Residues without data are shown in grey. (B) From the data presented in (A), the overall rotational correlation time τc, effective correlation times τe and generalized order parameters *S*^2^ for each ^15^N-^1^H vector were estimated by a Lipari-Szabo model-free analysis. The deduced τc was approximately 14.7 nsec. Meanwhile, theoretical τc values for a hypothetical dimer and a tetramer were calculated as follows. The two complex conformations were assumed to be spherical, and the radii of gyration were set to approximately 25 Å and 35 Å, referring to the crystal structures of the heterodimer and the heterotetramer. By applying these values of radii to the Stokes-Einstein-Debye equation, the rotational correlation times for the hypothetical dimer and tetramer complexes were estimated to be approximately 14.2 nsec and 38.8 nsec, respectively.

**Figure S8.**
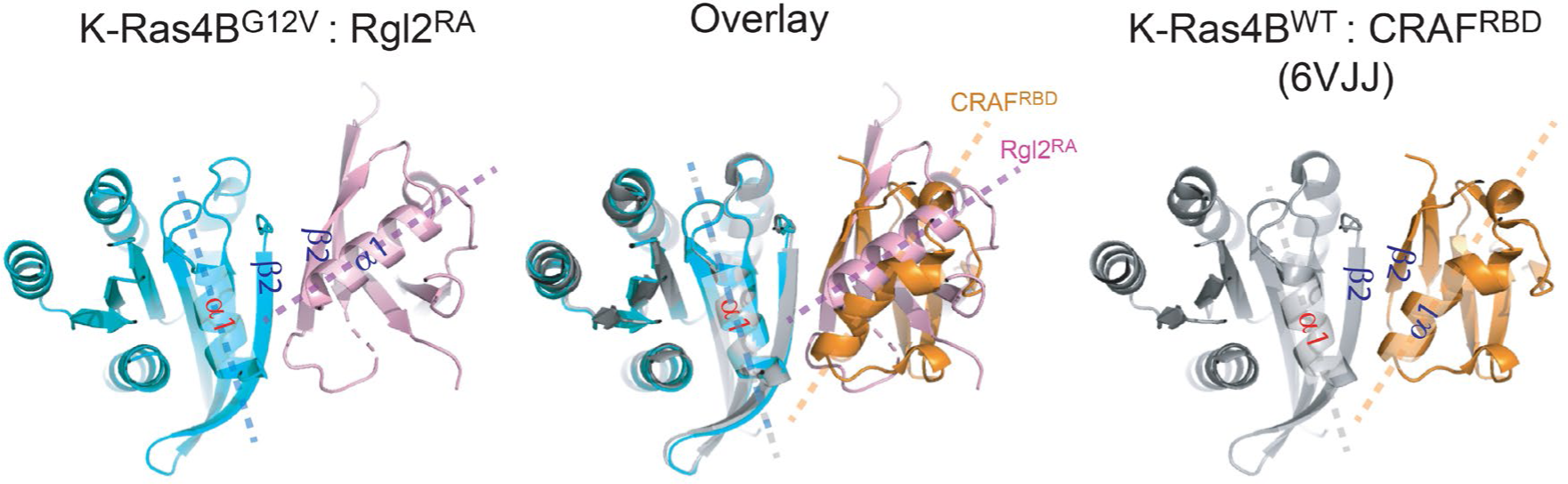
Comparison between KRas4B^G12V^:Rgl2^RA^ Switch I contact and KRas^wt^:CRAF^RBD^ complex. KRas4B^G12V^:Rgl2^RA^ complex formed through KRas4B Switch I region was compared with one of the representative Ras:RBD complexes, KRas4B^WT^:CRAF^RBD^ complex (PDB: 6VJJ). Both structures show β2 helices of KRas4B and RA/RBD run parallel to create the interface of the complex. Meanwhile, regarding the spatial arrangements of the α1 of Ras and α1 of RA/RBD, the axes of KRas4B^G12V^ α1 and Rgl2^RA^ α1 cross at a wider angle than the axes of KRas4B^WT^ α1 and CRAF^RBD^ α1 do. This feature is shared among Ras:RalGDS-family complexes (Eves *et al*., 2022).

**Figure S9.**
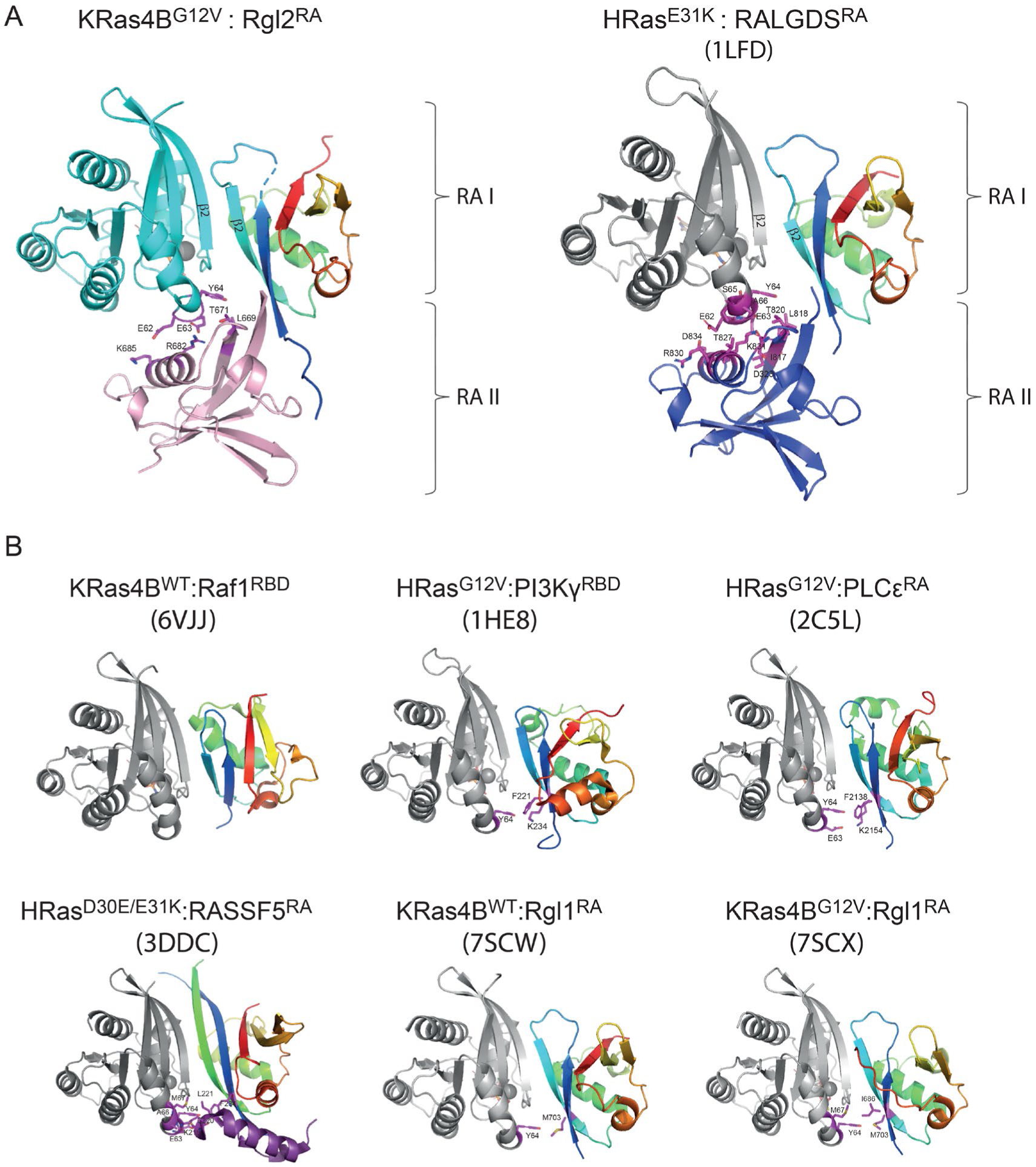
KRas4B^G12V^:Rgl2^RA^ and HRas^E31K^:RALGDS^RA^ complexes exhibit unique features among other Ras:effector complexes. The KRas4B^G12V^:Rgl2^RA^ complex was compared to representative Ras:effector complexes. The spectrum colour feature illustrates the orientation and ubiquitin super-fold status for one of the RA/RBD(s). When two RA/RBDs interact with one Ras molecule, the second RA/RBD is coloured pink (Rgl2^RA^) or blue (RALGDS^RA^). The KRas4B^G12V^ in the Rgl2^RA^ complex is in cyan, and other Ras molecules from previously published structures are in grey. The interacting residues involved in Ras Switch-II and RA/RBDs are depicted with magenta sticks and are labelled. (A) The two Ras:RalGEF complexes (KRas4B^G12V^:Rgl2^RA^ complex and HRas^E31K^:RALGDS^RA^ complex PDB: 1LFD) show similar structural features, including the orientation of the RAs, the Switch-I contact and the mode of usage of Switch-II. Two RAs (RA-I in the spectrum colour and RA-II in either pink or in blue) interact separately with a single Ras molecule at Switch-I and Switch-II regions. (B) Examples of Ras:effector complex crystal structures. KRas4B^WT^:CRAF^RBD^ complex (PDB: 6VJJ) shows that although Raf1^RBD^ has similar orientations of the ubiquitin fold structure to Rgl2^RA^-1, it does not utilise Ras Switch-II in complex formation. PI3Kγ^RBD^ (PDB:1HE8) and PLCε^RBD^ (PDB:2C5L) complexed with Ras utilise Switch-II region differently to Ras:RalGEF complexes despite a similar orientation of the ubiquitin fold structure of RA/RBDs. A single RBD interacts at both Switch-I and Switch-II regions of one Ras molecule. HRas^D30E/E31K^:RASSF5^RA^ complex (3DDC) shows yet another unique interaction where the additional α-helix at the N-terminal of the RA/RBD interacts with the Switch-II region of the Ras molecule, instead of the RA/RBD region itself. Rgl1 is one of the RalGEFs, as Rgl2 and RALGDS are. However, unlike Rgl2 or RALGDS, KRas4B^G12V^:Rgl1^RA^ complex (7SCX) and KRas4B^WT^:Rgl1^RA^ complex (7SCW) show one Rgl1^RA^ to interact with both Switch-I and Switch-II of one Ras molecule.

**Figure S10.**
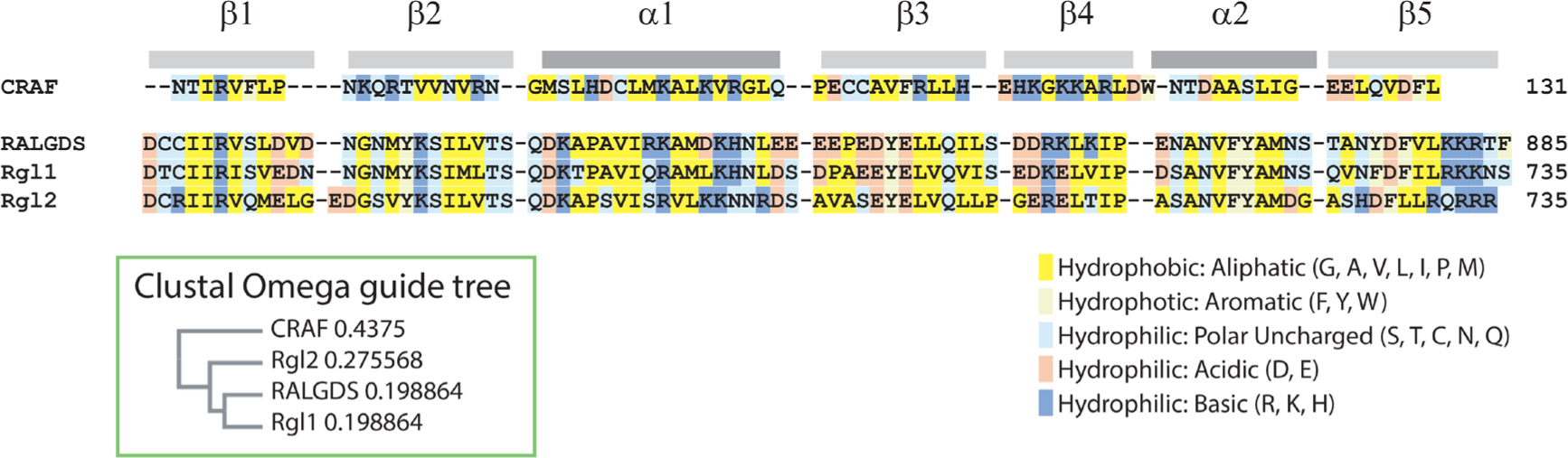
Rgl2^RA^ primary structure is highly similar to RalGEF^RA^ domains. Multiple sequence alignment of Ras binding domain (RBD) of human CRAF (residues 56-131) and Ras Association (RA) domains of human RalGEFs, RALGDS (residues 798-885), Rgl1 (residues 648- 735) and Rgl2 (residues 648-735), as defined by UniProt. The alignment, as well as the Clustal Omega guide tree (shown in a green-framed box)(Sievers et al., 2011), reveals that sequences are relatively divergent between CRAF^RBD^ and RalGEF family RA domains, although all RBD/RA domains share the common ubiquitin fold ββαββαβ structure. The RalGEF family RA domains share a high degree of sequence homology.

**Supplementary Figure S11.**
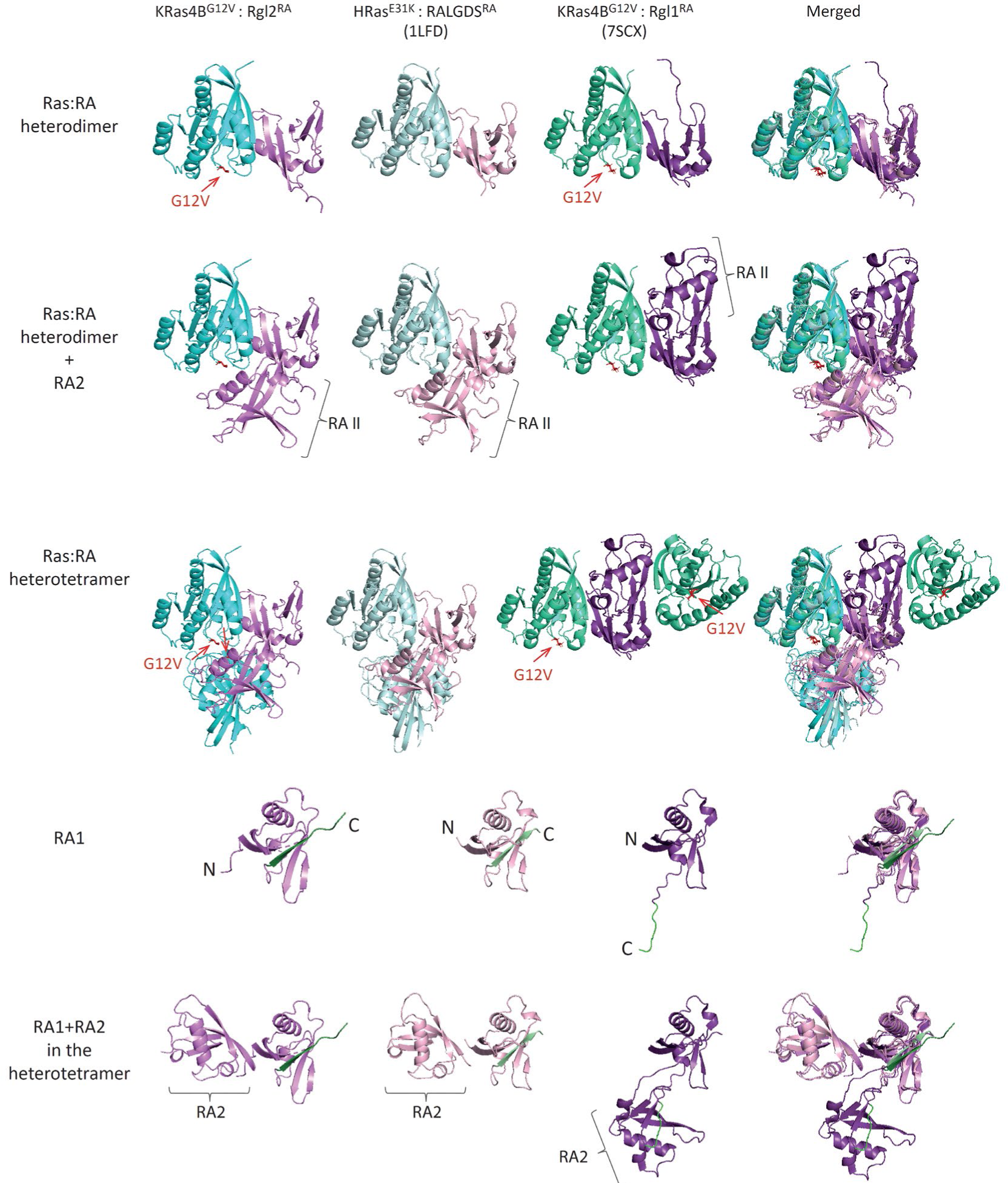
Heterotetramer formation in crystal structures of KRas4B^G12V^:Rgl2^RA^ and HRas^E31K^:RALGDS^RA^ complexes are distinct from the one in the KRas4B^G12V^:Rgl1^RA^ complex. Crystal structures of KRas4B^G12V^:Rgl2^RA^ (this study), HRas^E31K^:RALGDS^RA^ (PDB: 1LFD) and KRas4B^G12V^:Rgl1^RA^ (PDB: 7SCX) are compared. All the Ras:RA heterodimers, formed through the Ras Switch I region, show a highly similar structural arrangement (the top row). However, the way the second RA molecule interacts with the Ras:RA heterodimer in the KRas4B^G12V^:Rgl2^RA^ is shared only with the HRas^E31K^:RALGDS^RA^ complex (the second row). Consequently, the structural arrangements of the heterotetramers of KRas4B^G12V^:Rgl2^RA^ and the HRas^E31K^:RALGDS^RA^ crystal structures are distinct from the KRas4B^G12V^:Rgl1^RA^ heterotetramer (third row). RA domains of these structures also clearly show that Rgl2^RA^ and RALGDS^RA^ are structurally different from Rgl1RA, which interacts with the second Rgl1RA through the C-terminal end (two bottom rows). The C-termini of the first RAs are shown in green.

**Figure S12.**
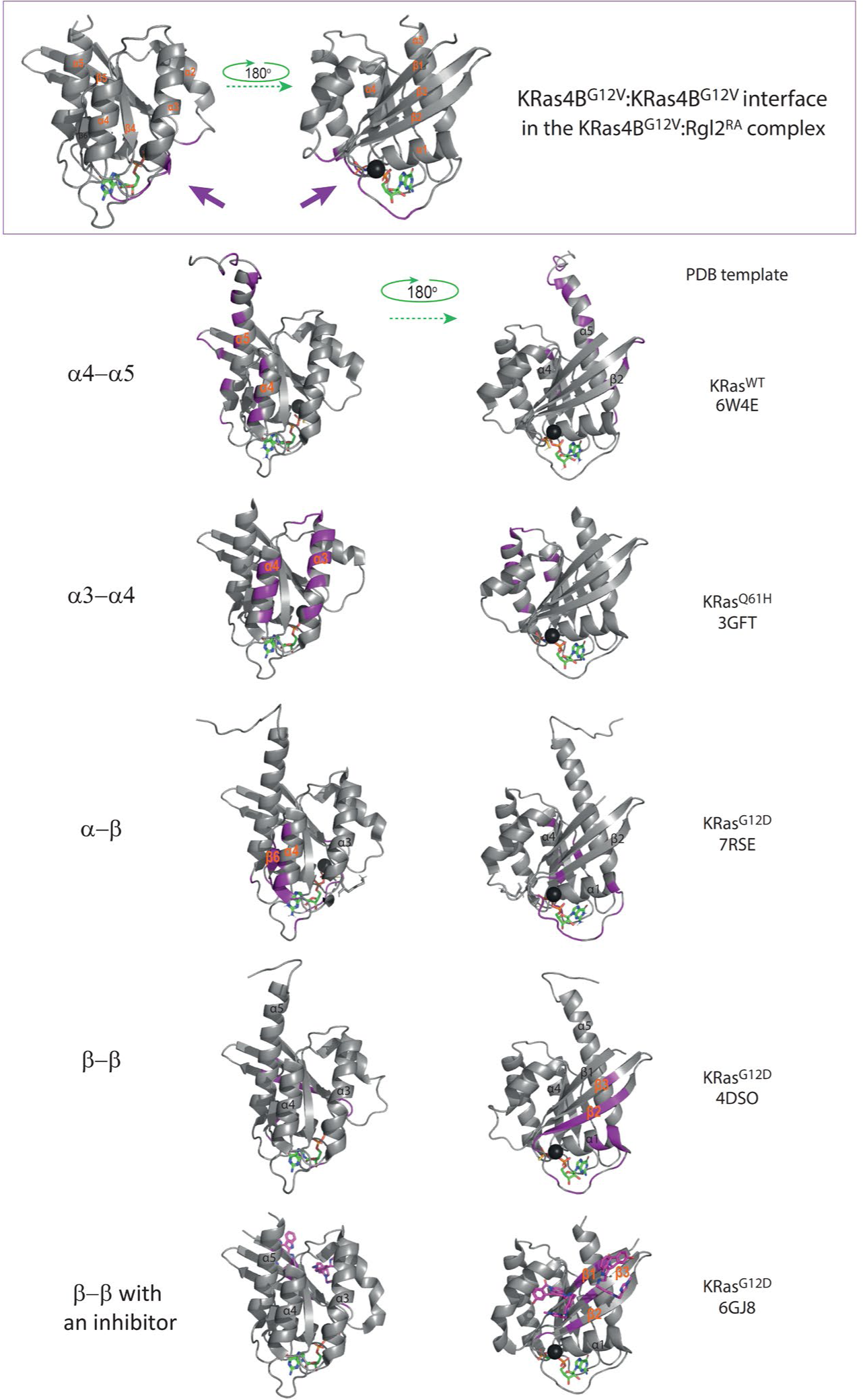
Various Ras:Ras interfaces proposed by experimental and computational prediction studies. The top panel shows the KRas4B^G12V^:KRas4B^G12V^ interface of the KRas4B^G12V^:Rgl2^RA^ heterotetramer complex (this study). The KRas4B^G12V^:KRas4B^G12V^ contacts (shaded in purple) are located in loop regions of Switch-I, Switch-II and α3. The KRas4B^G12V^:KRas4B^G12V^ interface is distinct from previously proposed dimerization interfaces, which can be classified into four categories based on the relevant structural elements; α5-α4, α4-α3, α-β and β-β. Example images for each category are shown. Single Ras molecules at the Ras:Ras interface are presented with the interacting residues highlighted in purple, and the involved α-helices and β-sheets are annotated. PDB IDs of the image templates are indicated next to each image. In the last example, the dimer formation was mediated by the β-β contact, aided by a small molecule inhibitor BI2852 (shown in purple). The Mg^2+^ is depicted as a dark-grey sphere, and the bound nucleotide is shown as a ball-and-stick model.

**Figure S13.**
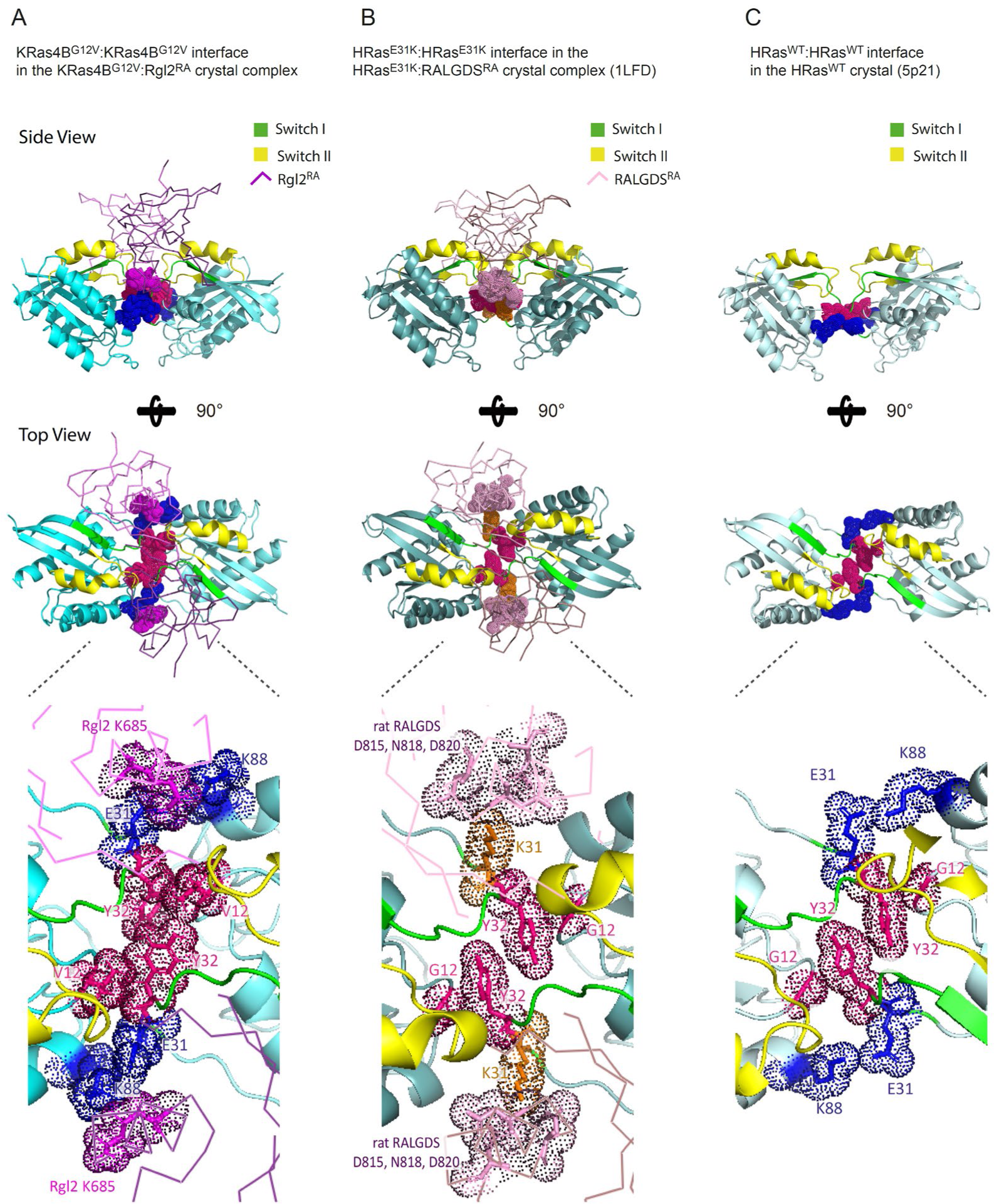
The non-canonical Ras:Ras contact seen in three crystal structures, KRas4B^G12V^:Rgl2^RA^ complex (this study), HRas^E31K^:RALGDS^RA^ complex (PDB ID 1LFD) and HRas^WT^:HRas^WT^ crystal structure (PDB ID 5P21) Neighboring Ras molecules are shown to reveal comparable spatial arrangements. The side view (top panel), top view (middle panel) and the blow-up images of the area Y32 of RAS (bottom panel) are shown. Switch I and Switch II regions are indicated in green and yellow. (A) KRas4B^G12V^:KRas4B^G12V^ in the KRas4B^G12V^:Rgl2^RA^ complex (this study). The two Rgl2^RA^ domains are shown as a ribbon in light magenta. (B) HRas^E31K^:HRas^E31K^ in the HRas^E31K^:RALGDS^RA^ complex (PDB ID 1LFD). The two RALGDS^RA^ domains are shown as a ribbon in light pink. (C) HRas^WT^:HRas^WT^ in the HRas^WT^ crystal (PDB ID 5P21). Spatial molecular arrangements around Y32 are similar in all the cases, but the oncogenic V12 provides a larger hydrophobic pocket, which may stabilise the KRas4B^G12V^:KRas4B^G12V^ interface of the KRas4B^G12V^:Rgl2^RA^ heterotetramer. (A and C) E31 of KRas4B^G12V^/ HRas^WT^ interact with KRas4B^G12V^/ HRas^WT^ K88. In (A), E31 of KRas4B^G12V^ also interacts with Rgl2^RA^ K685. (B) The hydrophobic pocket created by Y32 is not reinforced by G12, but K31 of HRas^E31K^ interacts with rat RALGDS^RA^ D815, N818 and D820 (the numbering was done according to Uniprot Q03386), facilitating the complex formation.

